# Root Remodeling Mechanisms and Salt Tolerance Trade-Offs: The Roles of HKT1, TMAC2, and TIP2;2 in Arabidopsis

**DOI:** 10.1101/2024.10.23.619678

**Authors:** N. O. Alshareef, V. J. Melino, N. Saber, A. De Rosa, E. Rey, J.Y. Wang, S. Al Babili, C. Byrt, M.A. Tester, M. M. Julkowska

**Affiliations:** Department of Biochemistry, Faculty of Science, King Abdulaziz University, Jeddah 80208, Saudi Arabia; Center for Desert Agriculture and Division of Biological and Environmental Sciences and Engineering, King Abdullah University of Science and Technology (KAUST), Thuwal, Saudi Arabia; Faculty of Science, Centre for Plant Science, School of Environmental and Life Sciences, University of Newcastle, Callaghan, NSW, Australia; Division of Plant Sciences, Research School of Biology, College of Science, Australian National University, Acton, Australian Capital Territory 2601, Australia; Boyce-Thompson Institute, Ithaca, New York, USA

## Abstract

Plant responses to salt stress involve complex processes integrating short- and long-term adaptations, including changes in ion transport, systemic signaling, root architecture, and biomass distribution. A key adaptive mechanism involves the regulation of sodium (Na^+^) and potassium (K^+^) ion transport via Class 1 HKT1 transporters, which reduce Na^+^ accumulation in shoots, thereby enhancing salinity tolerance but at the expense of lateral root development. In this study, we identified differential roles of TMAC2 in modulating ABA accumulation and lateral root development under salt stress in two distinct Arabidopsis genotypes, Col-0 and C24. Overexpression of TMAC2 in the Col-0 background increased ABA accumulation, resulting in reduced lateral root development, suggesting a positive feedback loop involving HKT1, TMAC2, and ABA signaling. In contrast, TMAC2 overexpression in C24 reduced ABA accumulation in lines overexpressing HKT1, indicating genotype-specific differences in the TMAC2-HKT1 interaction. Additionally, we observed that the co-expression of TMAC2 and HKT1 in Col-0 induced ABI4 and ABI5 transcription factors, which are known to mediate salt sensitivity. These findings reveal a regulatory network where TMAC2 and HKT1 modulate salt stress responses through genotype-dependent feedback mechanisms. Our results highlight the complexity of root remodeling under salt stress and the crucial role of genetic background in shaping these adaptive responses.

## Introduction

Plants exhibit a complex array of responses to salt stress, integrating immediate systemic responses (Choi et al. 2014; Evans et al. 2016), cell wall modifications (Feng et al. 2018; Zou et al. 2024), ion sequestration (Salazar et al. 2024; Møller et al. 2009), alterations of root system architecture (M. M. Julkowska et al. 2014, 2017), shift in biomass distribution between root and shoot (Ishka et al. 2024), and long-term adaptations that contribute to stress resilience and productivity (Awlia et al. 2021; Byrt et al. 2007; Munns et al. 2012). While the mechanisms behind the initial sensing and uptake of sodium chloride by plant roots remain elusive (van Zelm, Zhang, and Testerink 2020; Awlia et al. 2021), the strategies employed by plants to manage sodium distribution are comparatively well-documented (Ali et al. 2023; van Zelm, Zhang, and Testerink 2020). A key player in this management is the HKT1 transporter family, which regulates sodium (Na^+^) and potassium (K^+^) transport on a tissue level. Class 1 HKT1s preferentially transport Na^+^ out of the transpiration stream, thereby reducing the Na^+^ accumulation in the shoot, and thus limiting premature leaf senescence due to toxic Na^+^ accumulation in the photosynthetically active tissues (Davenport et al. 2007; Jaime-Pérez et al. 2017; Shamaya et al. 2017; Mäser et al. 2002).

Natural variation in the HKT1 promoter is associated with shoot Na^+^ content in dicots and monocots such as Arabidopsis and wheat (Rus et al. 2006; Baxter et al. 2010; Munns et al. 2012). When HKT1 was overexpressed in the stele shoot sodium content was reduced and improved salinity tolerance was observed (Møller et al. 2009). Interestingly, stelar overexpression of HKT1 also resulted in reduced lateral root development (M. M. Julkowska et al. 2017), resulting in reduced salt tolerance when salt stress was applied at an early developmental stage. This trade-off between salinity tolerance and lateral root development was hypothesized to be caused by increased sodium accumulation in the root xylem pericycle, which is also the tissue that gives rise to developing lateral root primordia (Dubrovsky et al. 2000; De Rybel et al. 2010; Goh et al. 2012). This duality of outcomes from stelar HKT1 expression highlights the need for a deeper understanding of the molecular controls orchestrating plant development under salt stress.

Lateral root development is a phenotypically plastic trait that was observed to change in response to nutrient and water availability. The molecular processes underlying lateral root development in response to water availability are well described in the context of hydropatterning or xero-branching (Bao et al. 2014; Orosa-Puente et al. 2018; Orman-Ligeza et al. 2018; Mehra et al. 2022; Johannes D. Scharwies et al. 2024). The temporary repression of lateral root growth in response to salt stress was previously ascribed to endodermal ABA signaling (Duan et al. 2013). However, the contribution of this response to overall salt tolerance and/or sodium exclusion is yet to be determined. While stele-specific overexpression of HKT1 led to repressed lateral root development under salt stress, supplementation of K^+^ alleviated the lateral root development (M. M. Julkowska et al. 2017). This observation provides an elegant tool to modulate lateral root development and sodium accumulation in root stele. The enhancer trap lines expressing HKT1 specifically in the pericycle cells (J2731; UAS_GAL4_:HKT1 in C24 background) have decreased shoot sodium content by 64% whereas expression in the vascular bundle and procambium excluding the pericycle cells (E2586 lines; UAS_GAL4_:HKT1 in the Col-0 background) have decreased shoot sodium content by only 37% (Møller et al. 2009).

Here, we used a transcriptomic approach to identify genes involved in root remodeling in response to salt stress downstream of HKT1. Functional characterization of loss of function mutants led us to identify two genes with salt-induced changes in lateral root development: a negative regulator of ABA (AtTMAC2) and an aquaporin isoform in the tonoplast intrinsic protein (TIP) subfamily (AtTIP2;2). Genetic manipulation of AtTMAC2 demonstrated an association between expression levels of AtTMAC2 and enhanced ABA accumulation in the root under salt stress. Our results suggest increased ABA is causal to impaired main and lateral root development, which is alleviated by AtTMAC2, a negative regulator of ABA response. We propose that the opposing effects of AtTMAC2 and HKT1 on the lateral root development are likely mediated by enhanced expression of ABI4, a DREB subfamily A-3 of ERF/AP2 transcription factor, and ABI5, a basic leucine transcription factors. Furthermore, we demonstrate that a vacuolar-localized aquaporin (AtTIP2;2) is a candidate for water and sodium transport, and inhibits lateral root development under salt stress. We demonstrate that AtTIP2;2 is expressed in the same tissue context as UAS_GAL4_:HKT1, where it regulates lateral root outgrowth independent of UAS_GAL4_:HKT1. Based on these results, we suggest that inhibition of lateral root development under salt stress is due to over-accumulation of sodium ions in the pericycle. In conclusion, this study sheds light on the regulation of root remodeling under salt stress and suggests that the process is associated with, but not directly regulated by, the Na+ transporter HKT1.

## Materials and Methods

### Plant Material and Growth Conditions

The two available lines with enhanced expression of HKT1 at its native site of expression (at the root stelar cells) were used in this study. These lines are E2586 UAS_GAL4_:HKT1 in the Col-0 background and J2731 UAS_GAL4_:HKT1 in the C24 background as described in (Møller et al. 2009). Seeds of the T-DNA insertion lines were obtained from the Arabidopsis Biological Resource Centre (ABRC) (**Supplementary Table S1**). The promoters and coding sequences of AtTIP2;2 and TMAC2 were cloned into GreenGate vectors (Lampropoulos et al. 2013). The promoters and coding sequences were cloned into pGGA and pGGC, respectively, with mutated BsaI sites using primers (**Supplementary Table S2**). Modular entry vectors were subsequently ligated with 35S / UBQ promoters and fluorophore tags (GFP or mCherry) into the destination vector (pGZ001) (Lampropoulos et al. 2013). All overexpression lines were generated by agrobacterium-mediated transformation of Arabidopsis thaliana Col-0, C24, E2586, or J2731. Oligos used in generating overexpression lines are listed in **Supplementary Table S2**. The primers to measure the gene expression in mutant lines are listed in **Supplementary Table S3**.

### Root Phenotyping

Salt-induced changes in root architecture were quantified according to (M. M. Julkowska et al. 2017). Four-day-old Arabidopsis seedlings were transferred to ½ MS square plates containing either 0 mM NaCl (control) or 75 mM NaCl (salt treatment). Roots were scanned every 2nd day until 10 days after treatment. RSA was quantified using the Smart Root (Lobet, Pagès, and Draye 2011) ImageJ plugin (version 1.53T).

### Sample Collection For RNA-Seq Experiment and Data Analysis

Four-day-old seedlings (UAS-HKT1 lines and background line) were transferred for 24 hours to ¼ MS plates supplemented with one of the following treatments: 0 mM NaCl, 75 mM NaCl, 30 mM KCl or 30 mM KCl with 75 mM NaCl. Roots from five-day-old seedlings were collected and combined from 150 to 200 seedlings for a single biological replicate and snap-frozen in liquid nitrogen. Total RNA was isolated using Trizol (Magdalena Julkowska 2018), and mRNA was enriched using Ambion Dynabeads (M. Julkowska 2020). mRNA was used for library preparation, and cDNA libraries were sequenced, producing 150 bp paired end-reads (NovaSeq6000, Novogene) with a data output average of 15 million reads per sample and Q30 of 85%.

Raw counts were normalized by TMM (edgeR package), and differentially expressed profiles were produced from three biological replicates by size factors (DESeq package). The expression of each mapped gene relative to Control conditions is listed in Supplementary Table S4. The normalized expression (tpmr) of each gene across individual samples is listed in Supplementary Table S5. The data was inspected for reproducibility using multi-dimensional scaling, and the differential expression was calculated using the DESeq package. We used the cutoff of -2 < log2(FC) > 2 and FDR p-value of 0.05 to identify differentially expressed genes for each condition x genotype combination using 0 mM NaCl 0 mM KCl as reference condition. We subsequently examined DEG shared between each genotype and condition and identified DEGs in response to treatment (75 mM NaCl, 30 mM KCl or 30 mM KCl and 75 mM NaCl) relative to control conditions (0 mM KCl and 0 mM NaCl), and HKT1 overexpression (E2586 vs E2586 UAS_GAL4_:HKT1 and J2731 vs J2731 UAS_GAL4_:HKT1). The genes showing response to both treatment and HKT1 were subsequently compared across individual background lines (Col-0 and C24) and salt stress treatments (75 mM NaCl and 30 mM KCl + 75 mM NaCl). The overlapping double-DEGs were inspected in further detail.

### Expression Analysis of Transgenic Lines

To determine the expression level of different transgenic lines, total RNA was extracted from the leaves of soil-grown Arabidopsis plants using Direct-zol RNA MiniPrep Plus (Zymo Research, CA, USA) following manufacturer’s instructions and quantified using a NanoDrop spectrophotometer. Total RNA (1 μg) was used to synthesize cDNA using iScript™ Reverse Transcription Supermix kit (Bio Rad) according to the manufacturer’s instructions. qPCR was performed using SsoAdvanced Universal SYBR Green Supermix (Bio-Rad, Hercules, CA, USA; 172-5270) on CFX96 real-time PCR machine (Bio-Rad) following standard protocol. The expression level of the target gene was calculated using the dCt method and normalized to the geometric mean of actin2 (AT3G18780) and EF1a (AT5G60390). Primers for qPCR are listed in Supplementary Table S3. Gene expression was calculated for three independent biological replicates and two technical replicates.

### Subcellular Localization Using Transient Co-Expression Assay

To determine the subcellular localization of AtTIP2;2 and AtTMAC2, each protein was transiently expressed in *Nicotiana benthamiana* leaf epidermal cells. Co-localization of AtTIP2;2 and AtTMAC2 with a marker protein was determined using signal correlation analysis based on confocal microscopy images. Three marker proteins were used in this study; plasma membrane protein 1;4 (PIP1;4) as a plasma membrane marker, AtVAMP711a as a vacuolar marker (Geldner et al. 2009), and Serrate as a nuclear marker, being expressed specifically in the nucleoplasm (Yang et al. 2006).

To obtain the expression vectors for protein transient expression, coding sequences of AtTIP2;2, AtTMAC2, AtPIP1;4, and VAMP711a were amplified from the cDNA of Col-0 Arabidopsis plants using Phusion High-Fidelity DNA Polymerase (New England Biolabs) and primers described in (**Supplementary Table S2**). Amplicons were purified and cloned into GreenGate entry vector pGGC (Addgene plasmid #48858; http://n2t.net/addgene:48858) to generate entry vectors (without stop codon). The entry vectors were subsequently cloned into the GreenGate (Lampropoulos et al. 2013) expression destination vector pGGZ001 (Addgene plasmid #48868; http://n2t.net/addgene:48868) using BsaI digestion and T4 DNA ligation (respectively M0202S and M0202S both from NEB). Correct ligation was confirmed by digestion with enzymes XmnI, BbsI, and BsmAI from NEB.

The 35S::AtTIP2;2-eGFP, 35S::AtTMAC2-eGFP, 35S:mcherry-AtPIP1;4, 35S::mCherry-VAMP711a, 35S::CFP-SERAT constructs were transformed into Agrobacterium tumefaciens strain GV3101 (Koncz and Schell 1986) using a heat-shock transformation protocol (Sparkes et al. 2006). Leaves from four-week-old *N. benthamiana* were inoculated with agrobacterium culture as follows: protein of interest (AtTIP2;2 or AtTMAC2), marker (AtPIP1;4 or AtVAMP711) and P19 (viral gene silencing suppressor to enhance transient expression efficiency (Voinnet et al. 2003). Co-localization was visualized 72 hr post-infiltration. Images were captured using a confocal laser scanning microscope (Stellaris, Leica). Images were analyzed using LAS X software (Leica). The eGFP excitation was at 488 nm, and the emission was between 500 and 530 nm, while the excitation for mCherry was at 561 nm, and the emission was at 580-630 nm, and for CFP 405 nm as excitation and 432-513 nm for emission.

### Localization of TIP2;2 and TMAC2 in Stable Transformants

To confirm localization in Arabidopsis under control of the native promoter, Arabidopsis transformants harboring either AtTIP2;2::AtTIP2;2-eGFP or AtTMAC2::TMAC2-eGFP were generated. Leaves and roots of ten days old Arabidopsis seedlings grown in ½ MS plates were harvested and visualized using confocal microscopy. Roots were stained with propidium iodide PI (10 mg.ml-1) for 1 min to better visualize the cellular structures of root cells. The PI was excited at 568 nm, and the emission was 585-610 nm. Images were captured using a confocal laser scanning microscope (Stellaris, Leica).

### Measurement of Na^+^ And K^+^ Content

Shoots and roots of Arabidopsis plants were collected after 21 days from transfer treatment plates (either 0 mM NaCl or 75 mM NaCl). Samples were then oven-dried at 75 ºC for about 3 days, digested by adding 5 mL of freshly prepared 1% (v/v) nitric acid (Sigma Aldrich), and incubated at 60°C for 2 days. The concentrations of sodium and potassium ions in the samples were determined using a flame photometer (model 425, Sherwood Scientific Ltd., UK), (Awlia 2019, Flame Photometry Protocol. https://dx.doi.org/10.17504/protocols.io.6t6here).

### ABA Quantification

Quantification of endogenous hormones was performed following the procedure in (J. Y. Wang et al. 2021). Up to 15 mg of freeze-dried ground root or shoot base tissues were spiked with internal standards D6-ABA (10 ng) and 1700 μL of methanol. The mixture was sonicated for 20 min in an ultrasonic bath (Branson 3510 ultrasonic bath), followed by centrifugation for 5 min at 14,000 × g at 4°C. The supernatant was collected and dried under a vacuum. The sample was re-dissolved in 120 μL of acetonitrile:water (25:75, v-v) and filtered through a 0.22 μm filter for LC–MS analysis. ABA was analyzed by LC-MS/MS using UHPLC-Triple-Stage Quadrupole Mass Spectrometer (Thermo ScientificTM AltisTM). Chromatographic separation was achieved on the Hypersil GOLD C18 Selectivity HPLC Columns (150 × 4.6 mm; 3 μm; Thermo ScientificTM) with mobile phases consisting of water (A) and acetonitrile (B), both containing 0.1% formic acid, and the following linear gradient (flow rate, 0.5 mL/min): 0–10 min, 15%–100 % B, followed by washing with 100 % B for 5 min and equilibration with 15 % B for 2 min. The injection volume was 10 μL, and the column temperature was maintained at 35 °C for each run. The MS parameters of Thermo ScientificTM AltisTM were as follows: negative mode, ion source of H-ESI, ion spray voltage of 3000 V, sheath gas of 40 arbitrary units, aux gas of 15 arbitrary units, sweep gas of 0 arbitrary units, ion transfer tube gas temperature of 350 °C, vaporizer temperature of 350 °C, collision energy of 20 eV, CID gas of 2 mTorr, and full width at half maximum (FWHM) 0.4 Da of Q1/Q3 mass. The characteristic Multiple Reaction Monitoring (MRM) transitions (precursor ion → product ion) were The characteristic MRM transitions (precursor ion → product ion) were 263.2 → 153.1, 263.3 → 204.1, 263.3 → 219.1 for ABA; 269.2 → 159.1 for D6-ABA.

### Screening for putative solute transport with high-throughput yeast-based assays

High throughput yeast microculture screens were used to test for putative transport of diverse key plant solutes (sodium, water, H_2_O_2_, boron and urea), associating growth enhancements or toxicity phenotypes to expression of foreign aquaporins (AQPs) in yeast cells in response to various treatments.

AtTIP2;1, AtTIP2;2 and AtTIP2;3 coding sequences with stop codon were cloned into Gateway-enabled expression vector pRS423-GPD-ccdB-ECFP (Simon and Gitler 2007) containing Uracil3 (URA3) yeast selection gene. Yeast expression vectors were transformed in respective yeast strains required for functional assays (described below), using the “Frozen-EZ yeast Transformation Kit II” (Zymo Research, Los Angeles, USA). Transformed colonies were grown in Yeast Nitrogen Base, YNB, media (Standard drop out, DO, -URA) and spotted (10μl) on agar YNB (DO -URA) selection plates for incubation at 30°C for 2 days, then stored at 4°C. Spotted plates (1/2 spot per tube) were used for the starting cultures of functional assays.

Water, hydrogen peroxide (H_2_O_2_), boron and urea functional assays were conducted as previously described in (Groszmann et al. 2023). Yeast lacking native aquaporin isoforms *aqy1aqy2* (null aqy1 aqy2; background Σ1278b; genotype: Mat α; leu2::hisG; trp1::hisG, his3::hisG; ura352 aqy1D::KanMX aqy2D::KanMX, (Tanghe et al. 2002) was used for the water/Freeze-thaw assay. The freeze-thaw assay exploits a property of yeast cells where increased freezing tolerance is conferred when the yeast is expressing functional water-permeable AQPs relative to yeast lacking functional water-permeable AQPs. The *aqy1aqy2* yeast was also used for a Boric Acid (BA) toxicity assay, which involved screening for enhanced AQP-associated sensitivity of yeast cells upon exposure to increasing BA treatment concentrations. H_2_O_2_ permeability was assessed using a reactive oxygen species (ROS) hypersensitive yeast strain, Δskn7 (null skn7; background BY4741 genotype: Mat α; his3Δ1 leu2Δ0 met15Δ0 ura3Δ0 ΔSKN7) (Bienert et al. 2007; Lee et al. 1999). In the ROS sensitivity assay yeast experienced enhanced toxicity responses if they were expressing AQPs that facilitated diffusion and accumulation of H_2_O_2_ in the cell. The Urea growth-based assay involved using *ynvwI* yeast (null dur3; background Σ23346c; genotype: Mat α, Δura3, Δdur3) containing a deletion of the DUR3 urea transporter (L.-H. Liu et al. 2003). In the Urea assay expression of urea-permeable AQPs in *ynvwI* yeast provided a growth advantage when exposed to media containing urea as the sole nitrogen source.

Our assay methodology framework was further adapted to establish a novel sodium toxicity screen using the salt sensitive yeast strain AXT3 (Δena1-4::HIS3 Δnha1::LEU2 Δnhx1::TRP1 (Quintero, Blatt, and Pardo 2000). AXT3 yeast cell survival was further compromised if they were expressing AtTIPs that facilitated the diffusion and accumulation of Na^+^ in the yeast cell, because this enhanced the toxicity response of the cell upon exposure to increasing NaCl treatments. AXT3 yeast expressing AtTIP2 isoforms and Empty vector control yeast were grown for 24 hours in 1 mL YNB(-URA) 10mM ethylene glycol-bis(β-aminoethyl ether)-N,N,N′,N′-tetraacetic acid (EGTA) pH6 media, at 30°C, shaking at 250rpm and diluted to 0.6×10^7^ cells/mL. 200μL microcultures of each AtTIP2/Empty vector were distributed in 96-well plates with 180μL of yeast and 200μL NaCl treatments: 0mM/water, 50mM, 100mM and 200mM NaCl.

Yeast microculture growth for all solute transport assays was monitored in 96-well plate using a SPECTROStar nano absorbance microplate reader (BMG Labtech, Germany) at 10–20-minute intervals over 42 to 60 hours. Data collection and processing for all solutes screened was conducted as described in (Groszmann et al. 2023). Briefly, growth curves were integrated using the natural log of OD_650_/initial OD_650_ (Ln(OD_t_/OD_i_) vs time) up to a time when the growth rate of the Untreated culture had declined to 5% of its maximum. Area Under the Curve (AUC), was calculated as a proxy that captured the potential growth characteristics affected, regardless of the treatment, in a single parameter. Statistical analyses were conducted using GraphPad PRISM Version 9 (San Diego, California, USA, www.graphpad.com). Error bars on graphs represent SEM. One-way ANOVAs and Post-Hoc Fisher’s LSD tests were conducted to test statistical difference of means between each AtTIP and Empty vector control. Asterisks were used to denote statistical significance *p<0.05, **p<0.01 and ****p<0.001. For each assay, 6 biological replicates were tested over 3 experimental runs.

## Results

### Identification Of *HKT1*-Dependent Patterns Underlying Reduced Lateral Root Development

To investigate the HKT1-related response to salt stress at the transcriptional level, we performed RNAseq on Arabidopsis seedlings from UAS_GAL4_:HKT1 enhancer trap lines in Col-0 and C24 backgrounds (Møller et al. 2009). Four-day-old seedlings were transferred to 0, 75 mM NaCl, 30 mM KCl, or 30 mM KCl with 75 mM NaCl. After 24h of treatment, root tissue was collected for RNA isolation and library preparation. Multidimensional scaling (**Fig. 1A**) revealed that individual replicates clustered together. Additionally, we observed that all samples from salt-treated plants (75 mM NaCl or 75 mM NaCl + 30 mM KCl) clustered separately from the control samples (0 mM NaCl or 30 mM KCl) on Dimension 1. Additionally, samples derived from Col-0 background (E25586, pink and black in **Fig. 1A**), or C24 background (J2731, purple and blue in **Fig. 1A**) clustered along Dimension 2. This pattern reveals that the effect of salt treatment and genetic background was the strongest on the observed transcriptional patterns. The presence of UAS_GAL4_:HKT1 resulted in a significant number of differentially expressed (DE) genes related to root developmental processes (**Fig. 1B**). The presence of salt (75 vs. 0 mM NaCl) resulted in DEGs related to anion transport and response to cadmium, both of which involve membrane transport genes and responses to charged ions (**Fig. 1C**). The simultaneous treatment of potassium and sodium chloride (30 mM KCl + 75 mM NaCl) resulted in a similar enrichment of genes in KEGG categories as by NaCl alone but with an increased number of genes and a greater significance of the response to starvation (**Fig. 1D**). These data support our earlier findings that KCl alleviates lateral root developmental defects caused by NaCl toxicity (M. M. Julkowska et al. 2017).

**Figure 1.**
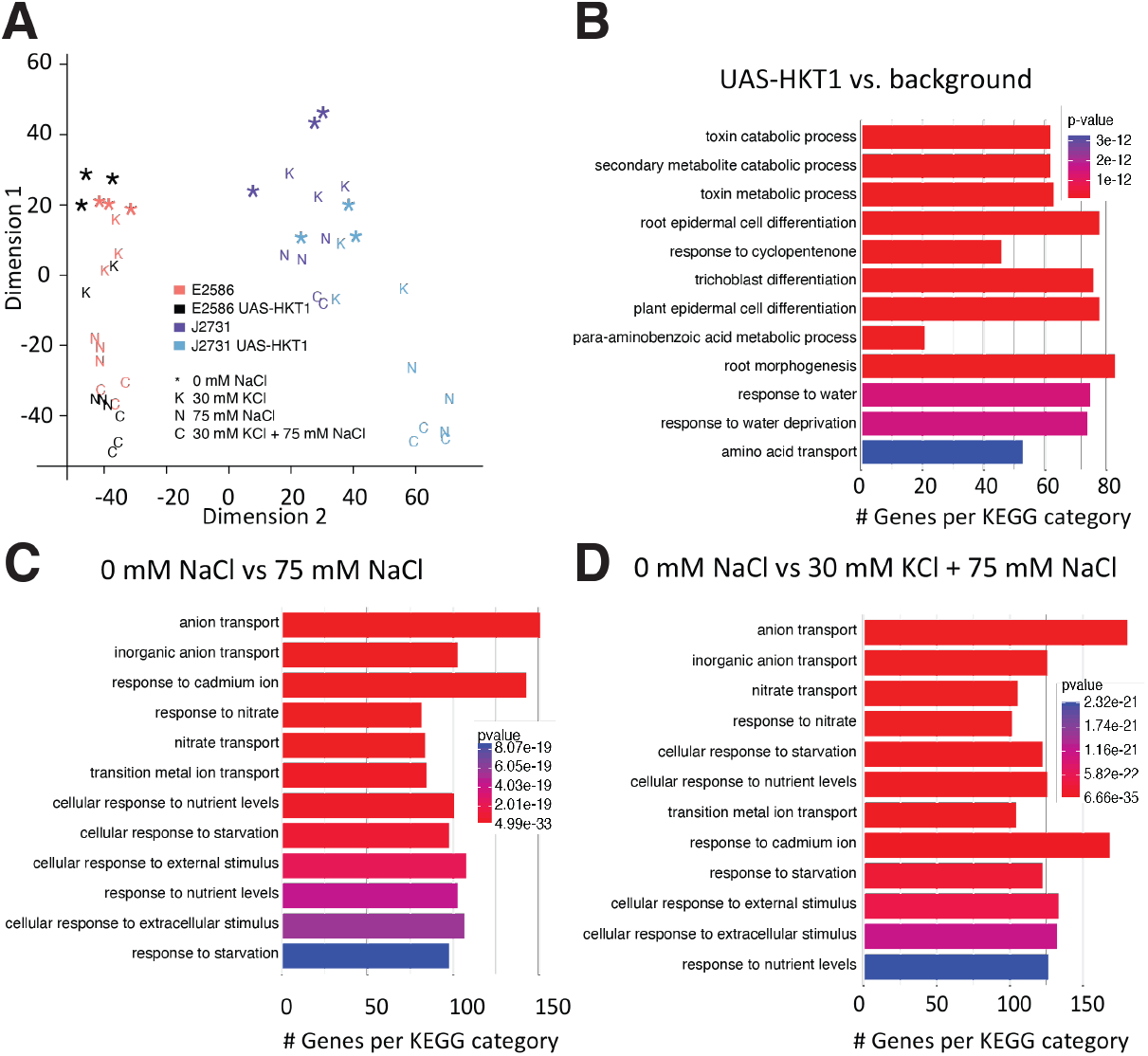
Pronounced effect of genetic background and treatment on gene expression in lines with tissue-specific overexpression of HKT1. **(A)** Principal component analysis of variance for each RNAseq sample grouped per treatment (*, 0 mM NaCl; K, 30 mM KCl; N, 75 mM NaCl and C, 30 mM KCl + 75 mM NaCl) and per genetic line of either the controls in Col-0 (E2586) or C24 (J2731) or with *UAS*_*GAL4*_:*HKT1* enhancer trap lines. The abundance of differentially expressed genes was evaluated per KEGG category for pair-wise comparisons **(B)** *UAS*_*GAL4*_:*HKT1* versus background; **(C)** control 0 mM versus stress at 75 mM NaCl and **(D)** control 0 mM NaCl versus supplemental K+ (30 mM KCl) during salt stress (75 mM NaCl).

To identify the genes downstream of UAS-HKT1 that may be involved in the repression of lateral root development under salt stress, we identified the genes induced by HKT1 overexpression in both Col-0 and C24 backgrounds (purple Venn, **Fig. 2A**). We identified three upregulated genes (AT3G02140 encoding TMAC2 - negative ABA regulator; AT5G05960 and AT3G18280 both encoding seed storage 2S albumin superfamily protein) and two downregulated genes (AT3G11340 encoding UDP-dependent glycosyl transferase 76B1; AT2G37060: NUCLEAR FACTOR Y, SUBUNIT B8) common to the presence of UAS_GAL4_:HKT1. Upon closer inspection, one of these genes, AtTMAC2 (AT3G02140), was up-regulated in salt (75 mM NaCl) conditions in the presence of UAS_GAL4_:HKT1 but only in the Col-0 background **(Fig. 2B**). AtTMAC2 is predicted to encode an ABI-5 (AFP4) binding protein and an ABA response element (ABRE) binding protein in the clade of bZIP transcription factors induced by ABA (Garcia et al. 2008; Lynch et al. 2017). Native expression of TMAC2 is predominantly found in the primary and lateral roots, and it was defined as a negative regulator of ABA and salinity responses. However, ABA data was not shown (Huang and Wu 2007). The salt treatment applied in the present study positively induced expression of TMAC2 in the C24 control, but UAS_GAL4_:HKT1 dampened this response, suggesting a negative feedback loop exists.

**Figure 2.**
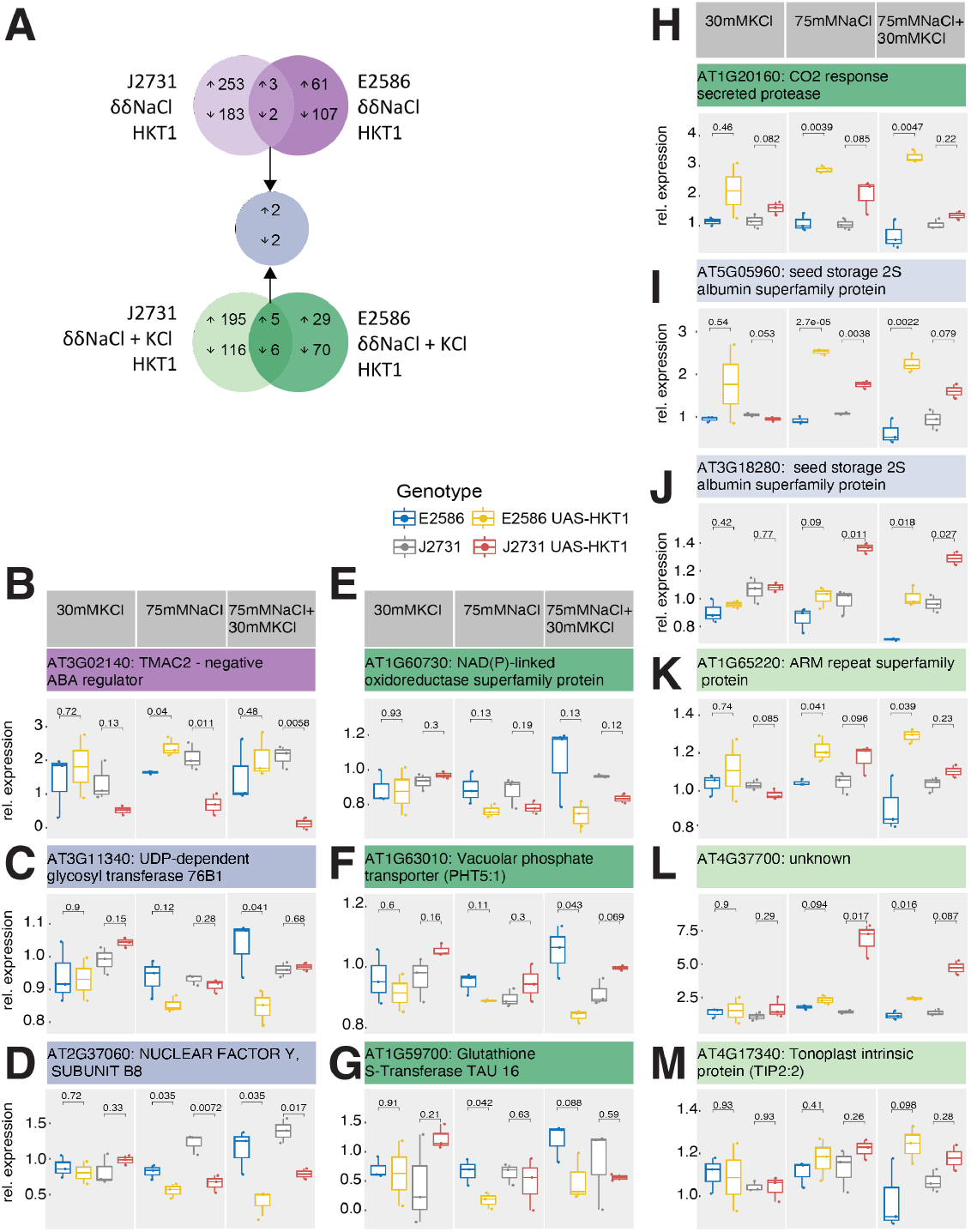
Identification of genes that exhibit similar regulation pattern in response to salt and HKT1 expression. **(A)** The differentially expressed genes were grouped based on their up- or down-regulation in either treatments (δNaCl for salt induced changes for 100 mM vs 0 mM NaCl, δ NaCl + KCl for salt induced changes for 100 mM NaCl + 30 mM KCl vs 30 mM KCl) or genetic backgrounds (δHKT1 J2731 for UAS-HKT1 in J2731 vs J2731; δHKT1 E2586 for UAS-HKT1 in E2586 vs E2586). The overexpression lines (UAS-HKT1) in both C24 (J2731) and Col-0 (E2586) backgrounds were compared to their respective backgrounds in response to NaCl (purple circles) and NaCl + KCl treatments (green circles). The genes shared between the genetic backgrounds and treatments were further inspected for their expression relative to control conditions across treatments, with **(B)** one transcript to show significant change in NaCl, **(C-D)** two transcripts to show significant reduction in response to both NaCl and NaCl + KCl treatment, **(E-H)** four transcripts to show significant reduction in response to NaCl + KCl, **(I-J)** two transcripts to show significant increase in response to both NaCl and NaCl + KCl treatments, and **(K-M)** three transcripts to show increase in response to NaCl + KCl treatment. The p-values listed above the individual comparisons between UAS-HKT1 and their background were calculated using T-test.

Common DEGs responsive to salt stress conditions (NaCl alone and NaCl + KCl) and HKT1 were identified (blue Venn, **Fig. 2A**). Two genes were specifically down-regulated, including a gene predicted to encode a UDP-dependent glycosyl transferase 76B1 (*At3g11340*, **Fig. 2C**) that may function in salicylic acid glucosyltransferase and a putative nuclear transcription factor Y subunit B-8 (*At2g37060*, **Fig. 2D**). There was no evidence for the other two components, NF-YA and NF-YB of the transcription factor Y, in their differential regulation in response to HKT1 and salt stress. Two genes involved in lipid transport from the bifunctional inhibitor/lipid-transfer protein 2s albumin superfamily (*At5g05960* and *At3g18280*) were significantly up-regulated in response to either salt stress conditions, but only in the presence of UAS-HKT1 (**Fig. 2I-J**). The protein encoded by *At5g05960* has a predicted plant lipid transfer domain also found in non-specific lipid transfer proteins involved in systemic acquired resistance and in the accumulation of cuticular wax and suberin (Finkina et al. 2016). These results suggest that while the phenotype of both UAS-HKT1 lines were very similar, their responses to both salt stress conditions differed severely in each background.

We examined the phenotypes of available mutants including the AT1G63010 - PHT5:1 (Vacuolar phosphate transporter, **Fig. S1**), AT3G11340 - UGT76B1 (UDP-dependent glycosyltransferase 76 B1, **Fig. S2**), and AT1G20160 - SBT5.2 (CO2 responsive secreted protease, **Fig. S3**). The majority of loss-of-function mutants of PHT5;1 showed increased lateral root emergence under salt stress compared to Col-0 (**Fig. S1, A-B**), whereas gain-of-function mutants showed decreased lateral root emergence (**Fig. S1, C**). Neither loss-nor gain-of-function mutants of UGT76B1 showed consistent differences with Col-0 in their lateral root development (**Fig. S2**). In contrast, their main root length showed significant differences under control and salt stress conditions (**Fig. S2**). Similarly, T-DNA insertion lines in SBT5.2 did not exhibit changes in lateral root development but showed increased main root length under control and salt stress conditions (**Fig. S3**). These results suggest that the DEG shared between two salt stress conditions and UAS_GAL4_::HKT1 lines play a role in root growth and lateral root development under both non-stress and salt stress conditions.

### The effect of TMAC2 on ABA accumulation depends on genetic background and HKT1 transcription

Previous reports of *AtTMAC2* (AT3G02140) demonstrated that although roots of the RNAi mutant (*tmac2)* were similar to that of the wildtype plant, overexpression of *AtTMAC2* (*35s::TMAC2*) resulted in shorter main roots, which were insensitive to ABA-mediated inhibition. The root phenotypic response of *35S::TMAC2* to salt stimuli was not described, nor were the effects on lateral root outgrowth with or without stimuli. Therefore, we explored the impact of TMAC2 on root architecture based on its significant alteration in expression in response to both salt and HKT1 (**Fig. 2B**). Interestingly, the expression was altered in two opposite directions in both backgrounds - its expression was enhanced by HKT1 in Col-0 background line, while in C24 background the expression of TMAC2 was repressed in the presence of high HKT1 expression (**Fig. 2B**). ABA is also known to inhibit lateral root outgrowth in response to salt stimuli mediated by the auxin-independent pathway (De Smet et al. 2003), thereby AtTMAC2 was a promising candidate for the salt-induced lateral root outgrowth inhibition observed here. TMAC2 was reported to be a negative regulator of ABA responses (Huang and Wu 2007), we have observed an ABA accumulation pattern that reflected the transcriptional changes (**Fig. 3A**). These results suggest that both HKT1 expression as well as genetic context affect the expression of TMAC2, and thereby also ABA levels.

**Figure 3.**
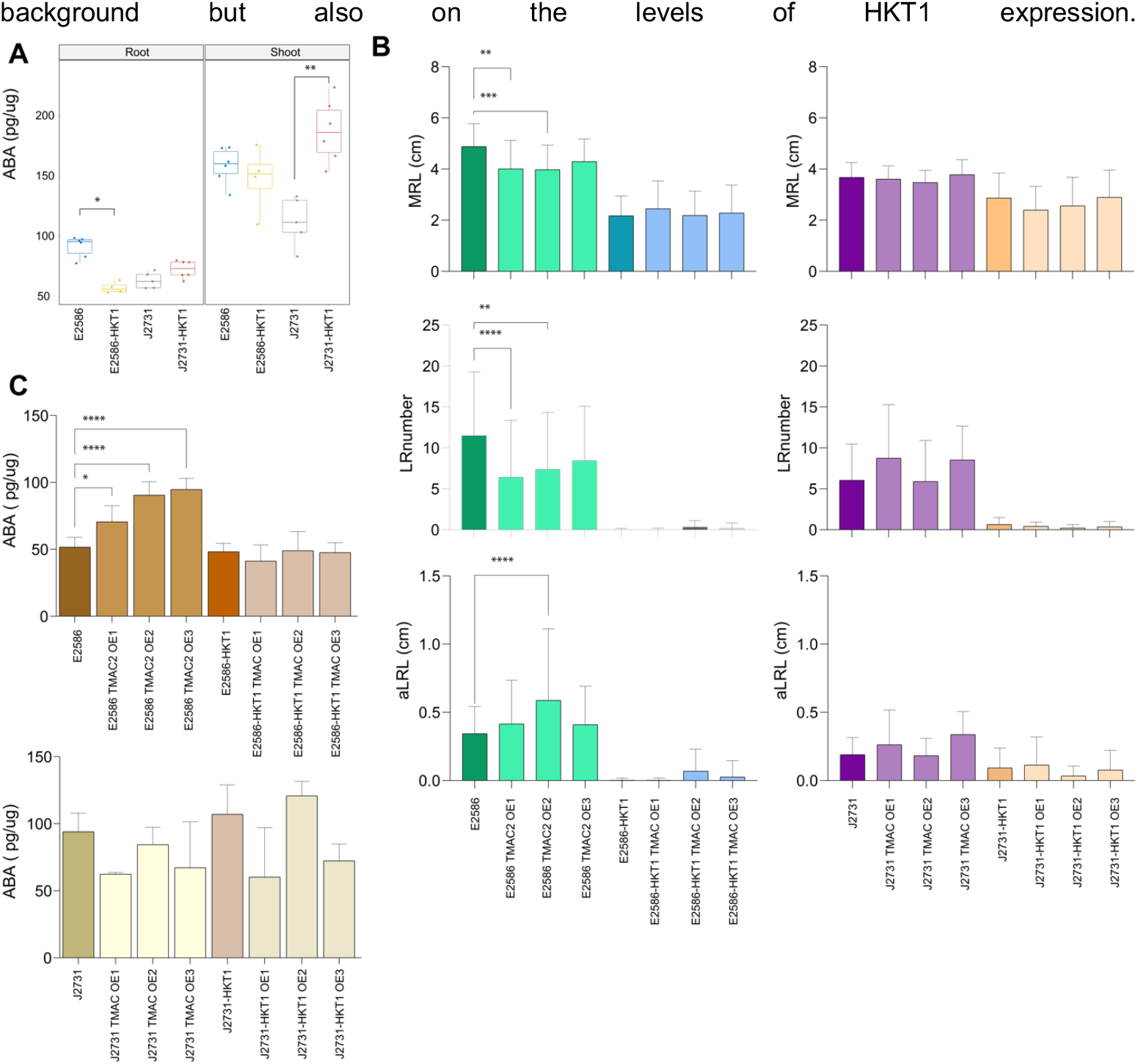
TMAC2 affects ABA accumulation and lateral root outgrowth. **(A)** The ABA accumulation in Col-0 (E2586) and C24 (J2731) seedlings with and without tissue specific overexpression of HKT1 was observed in 14 days old Arabidopsis seedlings, exposed to salt stress for 10 days. **(B)** Root system architecture, dissected into main root length (MRL), lateral root number (LR) and average lateral root length (aLRL) was determined for individual gain-of-function TMAC2 lines compared to their respective background linens of 13 days old plants exposed to salt stress (75 mM NaCl) for 9 days. **(C)** The ABA accumulation in the root tissue of gain-of-function TMAC2 lines and their background lines was measured from 18 days old seedlings, exposed to salt stress for 14 days. All of the measurements represent the average +/-standard error of at least 20 individual plants. The significant differences between individual mutant lines and their respective background lines were determined using one-way ANOVA test, with *, **, *** and **** indicating p-values below 0.05, 0.01, 0.001 and 0.0001 respectively.

To test whether lateral root development can be indeed altered through TMAC2 expression, we have generated constitutive expression of TMAC2 (*35S::AtTMAC2*, **Fig S4**) in four genetic backgrounds (Col-0 background with or without *UAS*_*GAL4*_:*HKT1*, and C24 background with and without *UAS*_*GAL4*_:*HKT1*). Although we anticipated that overexpression of TMAC2 would reduce lateral root outgrowth, we have only observed reduction in main root length and lateral root number in the Col-0 background (**Fig. 3B**) with no observable change in average lateral root length (**Fig. 3B**). No further decrease in main or lateral root number or length was observed either in C24 or in Col-0 *UAS*_*GAL4*_:*HKT1* (**Fig. 3B**). This could be caused by the fact that Col-0 *UAS*_*GAL4*_:*HKT1*, C24, and C24 *UAS*_*GAL4*_:*HKT1* lines were already severely impaired in lateral root development under salt stress conditions. These results suggest that increased TMAC2 expression is reducing the lateral root development exclusively in the Col-0 background in the scenario when HKT1 is not highly expressed.

In addition to the root phenotypes, we also measured ABA content in the roots and shoots of plate-grown plants (**Fig. 3C**). Overexpression of *AtTMAC2* in Col-0 (35S::AtTMAC2/E) led to an increased ABA content in the roots (**Fig. 3C**), but this increase was not observed in Col-0 *UAS*_*GAL4*_:*HKT1* line (**Fig. 3C**). No difference in ABA accumulation was observed in the shoot tissue of lines overexpressing TMAC2 (**Fig. S5**). In contrast, there was a trend of decreased ABA content in the C24 backgrounds with overexpression of HKT1 (**Fig. 3C**). These results suggest that HKT1 enhances ABA accumulation in C24 compared to Col-0 background and that overexpression of TMAC2 alleviates these changes. In the Col-0 background, the overexpression of TMAC2 enhances ABA accumulation but not in the presence of high HKT1 expression. In C24, however, overexpression of TMAC2 results in reduced ABA levels only in combination with HKT1. These results suggest that the role of TMAC2 in regulating ABA depends not only on the genetic background but also on the levels of HKT1 expression.

As TMAC2 was confirmed to localize into the cell nucleus (**Fig. 4A**), and the root phenotypes related to TMAC2 overexpression were only evident in the Col-0 background line, we hypothesized the existence of feedback regulation between TMAC2 and HKT1. We observed that HKT1 expression was enhanced in TMAC2 overexpression lines, even beyond the overexpression levels observed in Col-0 UAS_GAL4_::HKT1 background line (**Fig. 4B**). Moreover, the co-expression of both TMAC2 and HKT1 resulted in upregulation of transcription factors ABA Insensitive 4 (ABI4) and ABI5 (**Fig. 4C**). In contrast, overexpression of TMAC2 alone did not enhance the expression of either ABI4 or ABI5 (**Fig. 4C**). These results suggest that there is transcriptional feedback between HKT1 and TMAC2.

**Figure 4.**
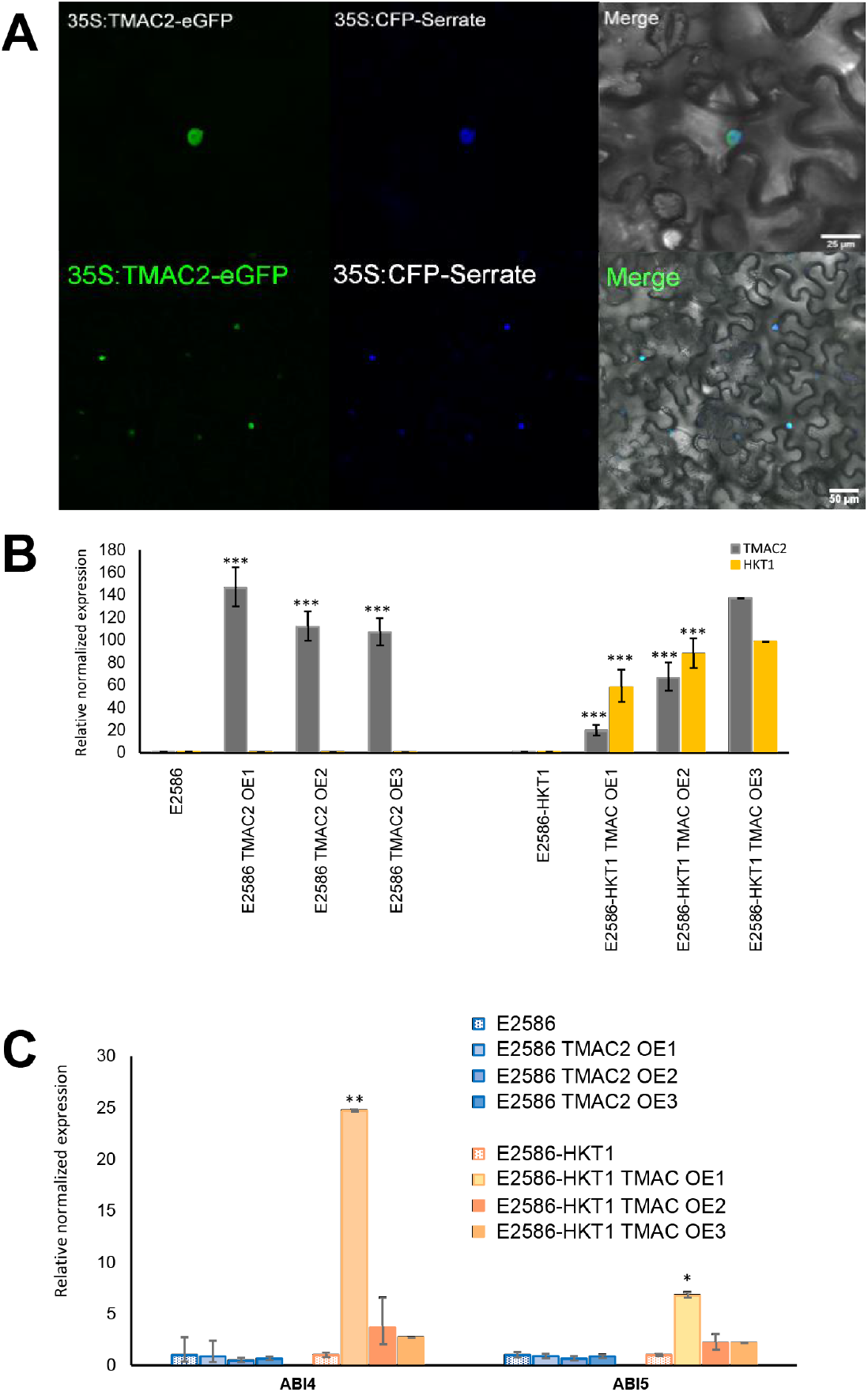
Feed-forward transcriptional regulation of TMAC2 and HKT1 through ABI4. **(A)** The subcellular localization of TMAC2 in transiently transformed *N. benthamiana* epidermal leaf cells. The cells were infiltrated with both 35S::TMAC2 construct and cell nucleus marker 35S::CFP-Serrate. The merged images illustrate co-localization of both GFP and CFP signals. **(B)** The abundance of both HKT1 and TMAC2 transcripts, **(C)** ABI4 and ABI5 transcripts relative to three housekeeping genes (Actin2, Ef-a and AtCACS (AT5G46630), measured in 18 days old seedlings of A. thaliana exposed to salt for 14 days. The graphs represent the average relative expression calculated from three biological replicates, while error bars represent standard error. The significant differences between individual mutant lines and their respective background lines were determined using one-way ANOVA test, with *, **, *** and **** indicating p-values below 0.05, 0.01, 0.001 and 0.0001 respectively.

### TIP2;2 regulates lateral root development in response to salt independent of HKT1

The RNAseq results revealed enhanced expression of TIP2;2 in HKT1 overexpression lines in both the Col-0 and C24 backgrounds, relative to wild type lines (**Fig. 2M**). TIP2;2 is an isoform in the tonoplast intrinsic protein subfamily of aquaporins, which are channels that can facilitate vacuolar membrane transport of water and other solutes (Loqué et al. 2005; Sudhakaran et al. 2021; Maurel et al. 1997). To investigate the function of the Arabidopsis TIP2;2 we evaluated the TIP2;2 T-DNA mutant in Col-0 background, as well as generated constitutive overexpression lines in Col-0 and C24 background with and without HKT1 overexpression (**Fig. 5**). The TIP2;2 loss-of-function mutant showed enhanced lateral root elongation (**Fig. 5A**), whereas the overexpression of TIP2;2 in either Col-0 or C24 background did not result in noticeable differences in root architecture under control and salt stress conditions (**Fig. 5B**), relative to wild type lines. Interestingly, overexpressing TIP2;2 in lines with tissue-specific overexpression of HKT1 did result in a further decrease of lateral root length (**Fig. 5C**), yet the effect was only significant in the C24 background. These results suggest that TIP2;2 has a negative effect on lateral root development under salt stress and that this effect is exacerbated by high HKT1 expression.

**Figure 5.**
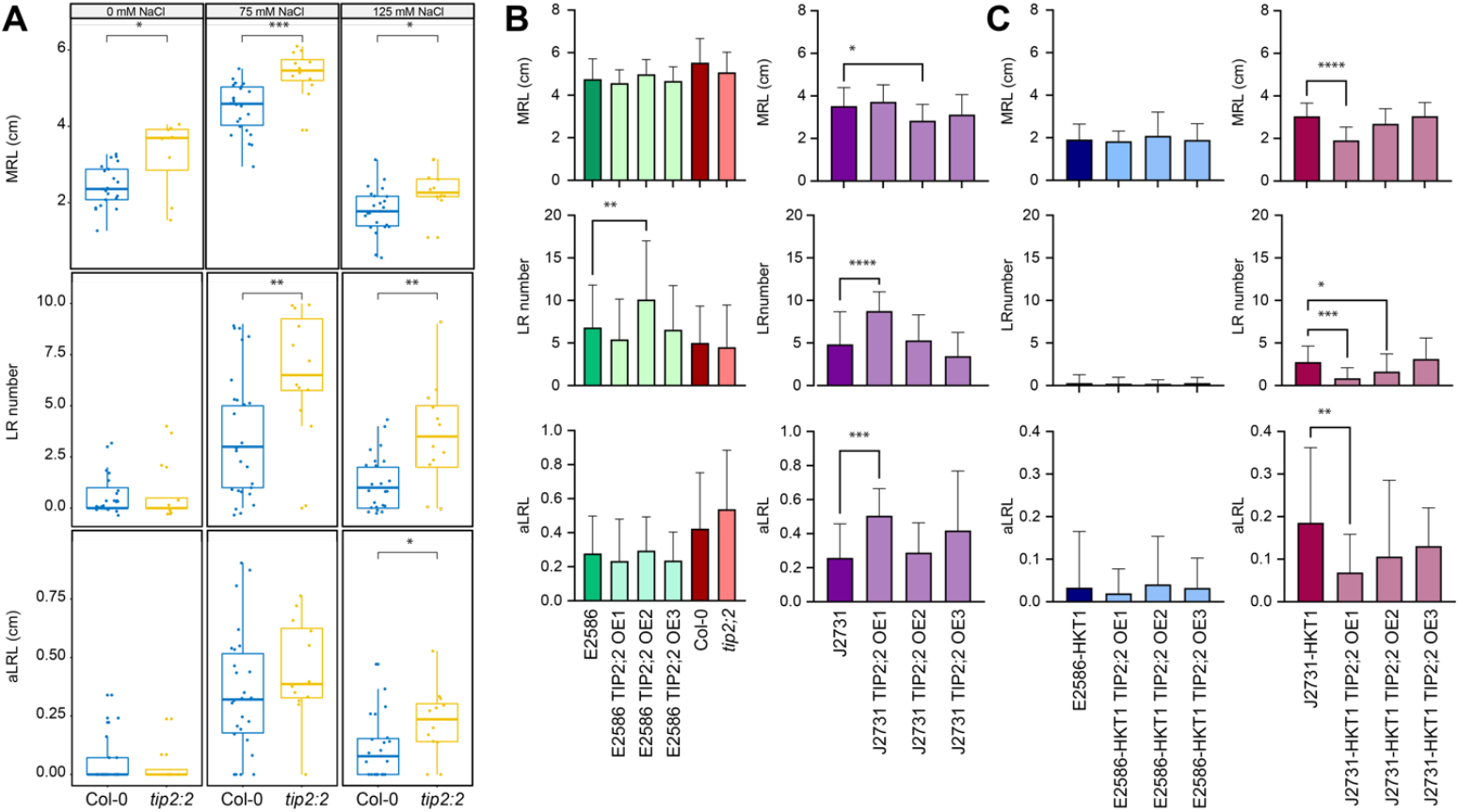
TIP2;2 is a negative regulator of lateral root development under salt stress conditions. **(A)** The T-DNA insertion line (*tip2;2*) was examined alongside Col-0 (wt) for salt-induced changes in root architecture. The root architecture was obtained from 8 days old seedling under non-stress conditions (0 mM NaCl) and 12 days old seedlings under salt stress conditions (75 and 125 mM NaCl). The seedlings were exposed to salt stress for 8 days. The box plots represent the interquartile range, while the thick line represents the trait mean. **(B)** The generated overexpression lines in E2586 and J2731 backgrounds, and **(C)** their respective tissue-specific HKT1 overexpression lines were examined for salt-induced changes in root architecture. The root architecture in this experiment was quantified of 13 days old seedlings exposed to 75 mM NaCl for 9 days. The bars represent the sample average and error bars represent standard error. The significant differences between individual mutant lines and their respective background lines were determined using one-way ANOVA test, with *, **, *** and **** indicating p-values below 0.05, 0.01, 0.001 and 0.0001 respectively.

To better understand the effect of TIP2;2 on lateral root development, we generated a translational fusion of AtTIP2;2 with GFP driven by the TIP2;2 native promoter and assessed it for sub-cellular localization (**Fig. 6**). We observed TIP2;2 to be highly expressed in the cortex of the elongation zone of the main root and mature lateral roots (**Fig. 6A - H**) and Arabidopsis leaf epidermal pavement cells (**Fig. 6I**). No TIP2;2 expression was observed in the lateral root primordia or early developing lateral roots (**Fig.6 B, C, E, H**), suggesting that its role is restricted to maturing tissues. TIP2;2 was observed in structures reminiscent of the plasma membrane, vacuole, and small vesicles (**Fig. 6J-P**). This sub-cellular localization was further confirmed through transient expression of 35S::AtTIP2;2::GFP in tobacco-infiltrated leaves (**Fig. 6Q-R**), demonstrating that AtTIP2;2 can localize to both the vacuolar and the plasma membranes. These results confirm the tonoplast localization of TIP2;2 and identify maturing roots and leaves as the primary tissues affected by TIP2;2 expression.

**Figure 6.**
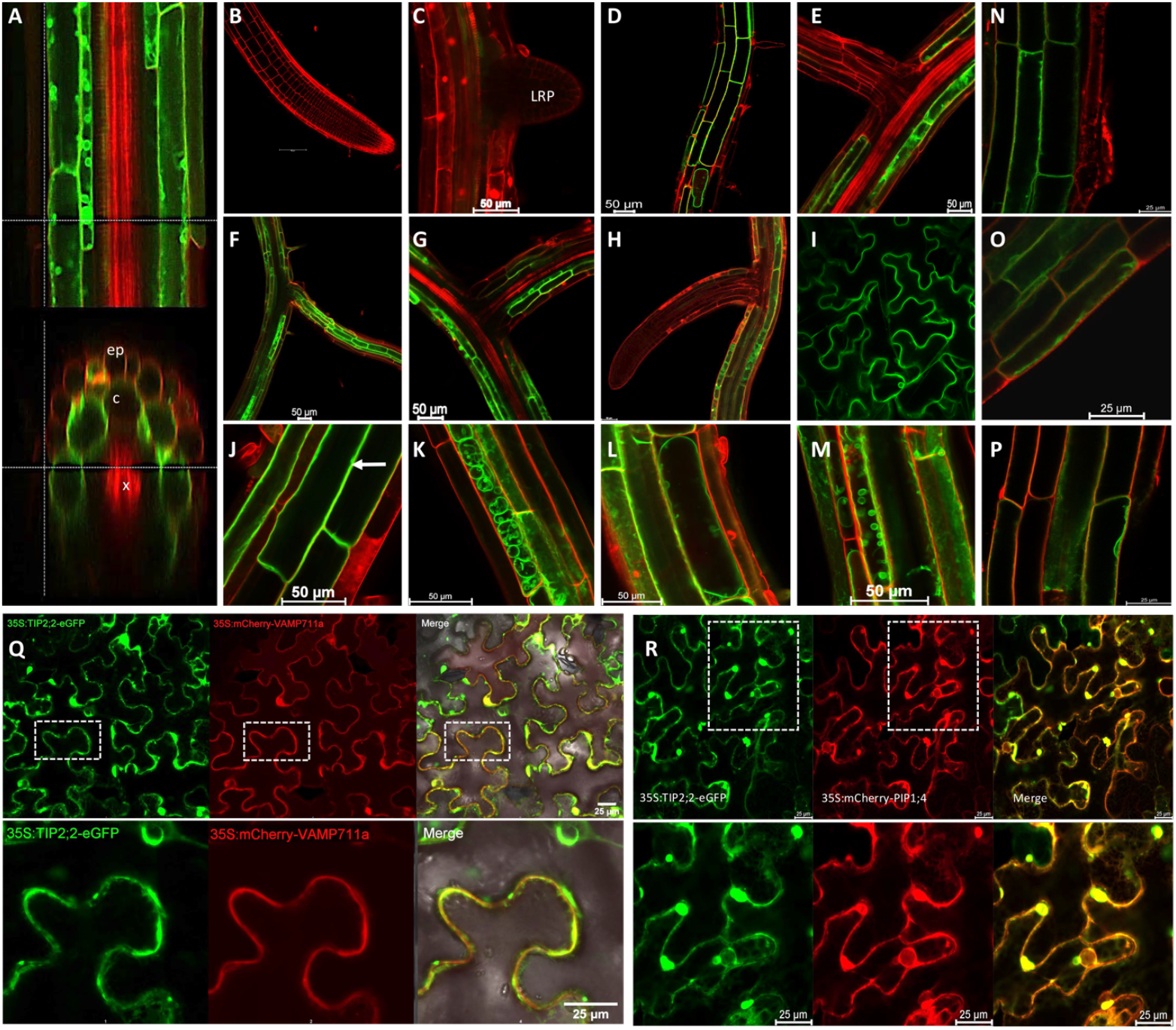
TIP2;2 is expressed in the cortex layer from the elongation zone onward and localizes to both tonoplast and plasma membranes. Ten days old Arabidopsis seedlings transformed with pTIP2;2::TIP2;2::GFP construct were additionally stained with propidium iodide and explored for tissue-specific expression and sub-cellular localization of TIP2;2. **(A)** The optical cross-section of the root elongation zone with epidermis (ep), cortex (c) and xylem (x) indicated on the image. Further the expression was investigated in **(B)** Main Root tip, **(C)** newly developing lateral root primordia (LRP), **(D)** maturation zone of the main root, **(E)** base of the developing lateral root, and **(F-H)** fully developed lateral root, **(I)** leaf epidermal pavement cells. We additionally observed various sub-cellular localization of TIP2;2 pointing towards **(J)** plasma membrane localization, **(K)** prevacuolar vesicle localization and **(L)** tonoplast localization, and **(M)** smaller vesicle localization. The additional staining performed with the membrane dye (FM4-64) was performed for **(N-O)** 5 minutes to confirm plasma membrane localization, and **(P)** 45 minutes to confirm vacuolar localization. The transient co-expression of p35s::TIP2;2::GFP in *N. benthamiana* epidermal cells **(Q)** in combination with vacuolar marker p35S::VAMP11a::mCherry and **(R)** in combination with plasma membrane marker p35S::PIP1;4::mCherry showed co-localization of TIP2;2 with both of the markers.

To investigate TIP2;2 permeability we progressed heterologous yeast-based microculture assays. Two AtTIP2 homologs from within the TIP subfamily, AtTIP2;1 (At3g16240) and AtTIP2;3 (At5g4750) were included in the assays exploring AtTIP2;2 permeability (**Fig. 7)**. High-throughput growth- and toxicity-based yeast assays described in Groszmann et al. (2023) were used to check whether these TIPs were permeable to water, urea, hydrogen peroxide and boric acid. This approach was further adapted to develop a novel sodium toxicity screen to check for putative sodium transport in TIP-expressing yeast cells, using a salt-sensitive yeast mutant strain AXT3 (Quintero, Blatt, and Pardo 2000). Yeast growth curves, Ln(OD/ODi) vs. time (**Fig. S10**) and Relative Area under the curve (AUC) measurements (**Fig. 7**) revealed that NaCl sensitive AXT3 yeast expressing AtTIP2;2-exhibited reduced growth compared to Empty vector control yeast upon exposure to 50 mM, 100 mM and 200 mM NaCl treatments, indicating that TIP2;2 is a candidate for facilitating Na^+^ transport into the yeast cells and causing a toxicity response. AtTIP2;2 was also observed to transport water, urea, hydrogen peroxide, and boric acid (**Fig. 7B-E, Fig. S10**). AtTIP2;1 and AtTIP2;2 were also observed to transport sodium, urea, and boric acid (**Fig. 7**), and TIP2;3 is was not identified as candidate for water and hydrogen peroxide transport, unlike the other two TIP2 isoforms. These results reveal that there is variation in the permeability of TIP2s to different plant solutes, and indicate that AtTIP2;2 and AtTIP2;1 have similar solute transport profiles, whereas AtTIP2;3 differed.

**Figure 7.**
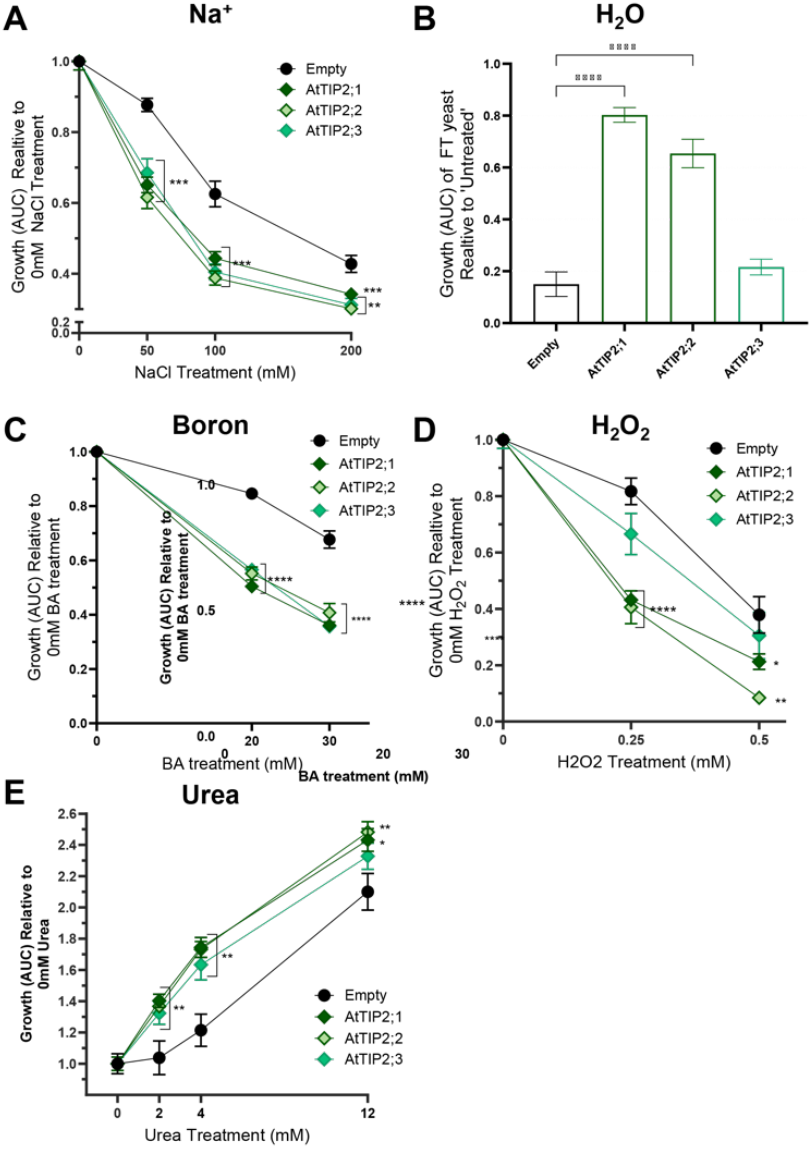
Solute transport profile phenotypes for AtTIP2;2 and its closest homologs, AtTIP2;1 and TIP2;3. High-throughput yeast screens were conducted to test for (A) Sodium, (B) Water, (C) Boron, (D) Hydrogen peroxide (H_2_O_2_) and (E) Urea permeability. Growth phenotypes were derived following (Groszmann et al. 2023) by calculating the Area Under the curve (AUC) from yeast OD_650_ vs time graphs for each culture exposed to various treatments. Sodium, Boron and H_2_O_2_ are toxicity-based screens, with decreased yeast growth/increased toxicity phenotypes of positive candidate TIP-expressing yeast cells, compared to an Empty vector control, indicative of TIP-mediated solute transport. Water transport assay (Freeze-thaw screen) attributes yeast survivorship enhancements of positive candidate TIP-containing yeast cells compared to Empty vector negative controls, reflective of respective levels of water-efflux/transport across membranes upon freezing and thawing treatments. The Urea growth-based assay enables comparison of growth enhancements of yeast grown in low-nitrogen treatments, where positive candidate TIP-expressing yeast displayed enhanced growth compared to Empty vector control, indicative of TIP-mediated urea transport. Asterisks denote One-Way ANOVA results comparing Treated yeast growth against Empty vector; *p<0.05, **p<0.01 and ***p<0.001. N=6, Error bars=SE. AtTIP2;1 and AtTIP2;2 were both identified as putative candidates for transport of sodium, water, boron, H_2_O_2_ and urea, resulting in enhancements in toxicity or growth/survivorship phenotypes in the respective assays. AtTIP2;3 was identified as a positive candidate for sodium, boron and urea transport, but not for water and H_2_O_2_.

To evaluate the function of TIP2;2 *in planta* we tested ion accumulation in TIP2;2 loss-of-function, overexpression and control lines. Overexpression of TIP2;2 did not result in enhanced sodium accumulation by itself but resulted in enhanced root Na^+^ content in the Col-0 line that had high HKT1 expression (**Fig. 8a**). We observed increased shoot K^+^ accumulation in TIP2;2 overexpression lines in Col-0 *UAS*_*GAL4*_:*HKT1* (**Fig. 8c**), while root K+ accumulation remained unaltered by TIP2;2 overexpression (**Fig. 8d**). No significant effect on ion accumulation was observed in the C24 background, independent of HKT1 expression (**Fig. S8**). These results indicate that AtTIP2;2 may have unique transport functions depending on the organ, and may enhance vacuolar storage of Na^+^ in the root while promoting vacuolar K^+^ retention in the shoot. The enhanced Na^+^ and K^+^ accumulation demonstrated here shows a direct requirement for *UAS*_*GAL4*_:*HKT1* and its dependency on genetic background.

**Figure 8.**
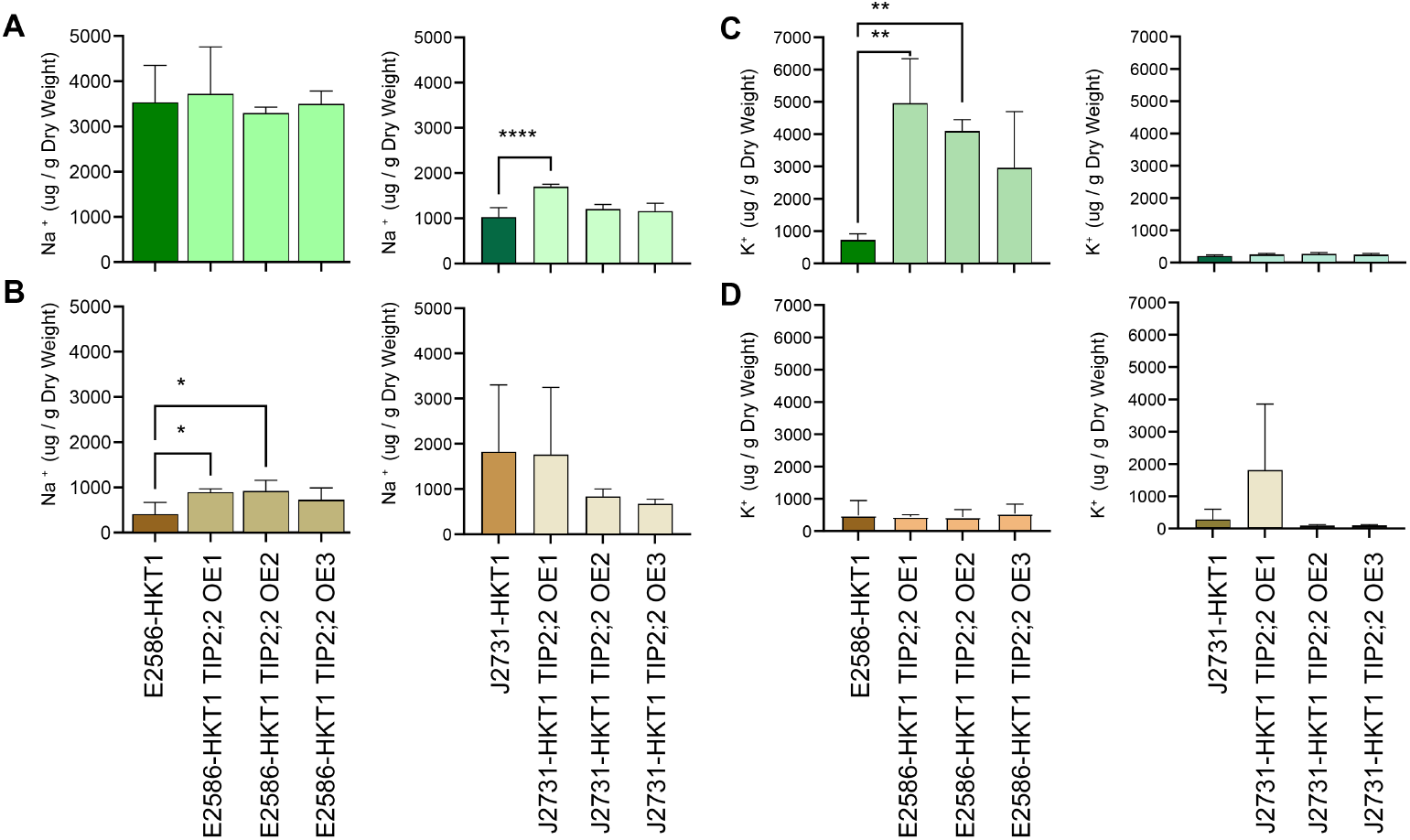
TIP2;2 contributes to sodium compartmentalization in the root. Four days old Arabidopsis seedlings from lines with tissue-specific HKT1 overexpression in Col-0 (E2586) and C24 (J2731) backgrounds with and without TIP2;2 overexpression were exposed to salt stress (75 mM NaCl) for 21 days. The sodium (Na^+^) accumulation was measured in seedling’s **(A)** shoot and **(B)** root tissue, as well as potassium (K^+^) accumulation was measured in seedling’s **(C)** shoot and **(D)** root. The bars represent the mean value calculated from at least 20 seedlings, and the error bars represent standard error. The significant differences between individual mutant lines and their respective background lines were determined using one-way ANOVA test, with *, **, *** and **** indicating p-values below 0.05, 0.01, 0.001 and 0.0001 respectively.

## Discussion

HKT1 is well known to contribute to shoot Na^+^ exclusion through retrieval of Na^+^ from the xylem, resulting in increased salt tolerance in a wide range of plant species (H. Song et al. 2023; Byrt et al. 2007; Munns et al. 2012; Jaime-Pérez et al. 2017; Rus et al. 2006). While root system architecture undergoes active reprogramming in response to salt stress (M. M. Julkowska et al. 2014), there is a trade-off between sodium exclusion and lateral root development that is regulated by HKT1 (Møller et al. 2009; M. M. Julkowska et al. 2017). Our running hypothesis for this trade-off is that the expression domain of HKT1 overlaps with the site of lateral root initiation, which is the xylem pole pericycle (Parizot et al. 2008; De Smet et al. 2007; Casimiro et al. 2003; Ramakrishna et al. 2019). Plants exposed to salt stress during their early development are impaired in their salt tolerance by high levels of HKT1 (M. M. Julkowska et al. 2017). This increases Na^+^ accumulation in the lateral root primordia, severely impairing lateral root emergence and seedling vigor (M. M. Julkowska et al. 2017). This effect can be partially rescued with additional K+ to the media (M. M. Julkowska et al. 2017). Here, we investigated the transcriptional responses to salt and high HKT1 expression in Col-0 and C24 backgrounds. Our results revealed that the genetic background and/or specific tissue at which HKT1 is expressed play major roles in eliciting transcriptional responses to salt stress (**Fig. 1A**). Previous studies in various organisms revealed the importance of promoter-enhancer architecture (Vihervaara et al. 2017; Yu et al. 2024; Das and Bansal 2019) as well as complex regulatory networks (Liang Song et al. 2016; Bjornson, Dandekar, and Dehesh 2016; Barah et al. 2016; Van den Broeck et al. 2017) that differ across genetic backgrounds and exhibit tissue-specific context. With the currently available resources we are not able to distinguish whether the major factor of transcriptional response differentiation between the studied lines is the specific overexpression site of HKT1 (pericycle specific overexpression in C24 background and vascular bundle and procambrium-specific overexpression in Col-0 background), or the genetic context (Col-0 vs C24). In the future, the studies of transcriptional responses across diversity panels (Ma et al. 2021; Pignon et al. 2021; Lin et al. 2022; Han et al. 2022) will help us further develop models underlying these complex networks and specific conditions for regulating stress-induced gene expression.

Although we have identified only 12 genes with altered expression by both HKT1 and salt stress in both lines, we did identify KEGG categories that were enriched in lines overexpressing HKT1 (**Fig. 1B**). One of the most surprising categories was the reprogramming of the epidermal cells, including the trichoblasts. Previous studies did observe that root epidermis undergoes significant changes in response to salt stress, including the severe reduction of root hair development (Cuin and Shabala 2007; Y. Wang et al. 2008; Arif, Islam, and Robin 2019; Y. Song et al. 2022; Ibeas et al. 2024). While the initial study of cell-type specific overexpression of HKT1 did observe slightly increased Na+ accumulation in the epidermal cell layers, its magnitude was far lower than other cell layers, such as xylem parenchyma, endodermis, and cortex (Møller et al. 2009).

We were able to validate the contribution to lateral root development of two out of 12 genes identified within this study. While we examined available mutant lines for their phenotypes regarding lateral root development, we might have missed other important contributions of these genes to plant development under salt stress, such as root hair development, as mentioned above. The loss-of-function mutants we studied in this study (**Fig. S1-S3**) did not exhibit significant root architecture phenotype due to either weak loss-of-function or gene redundancy. For example, the vacuolar phosphate transporter identified here (**Fig. 2F, AT1G63010**) might be involved in modulating response to salt stress in a HKT1-dependent manner, as salt is known to restrict phosphate availability and alter the overall responses (Kawa et al. 2016; Ibeas et al. 2024; Miura et al. 2011). Further investigations of these genes throughout gain-of-function and compound loss-of-function mutants will reveal their further contributions to root development under salinity stress, as well as their contributions to overall salinity tolerance.

The mechanisms underlying salinity reducing lateral root outgrowth were previously described to occur through prolonged spikes in endodermal ABA signaling that resulted in lateral root quiescence (Duan et al. 2013). Here, we demonstrated that *TMAC2* contributes to increased ABA accumulation in the Col-0 background (**Fig. 3C**) resulting in reduced lateral root development (**Fig. 3B**). As this TMAC2-dependent upregulation of ABA was not observed in UAS_GAL4_::HKT1 Col-0 line (**Fig. 3C**), we hypothesized that there might be a regulatory feedback loop between HKT1 and TMAC2. We have observed that overexpression of TMAC2 indeed resulted in increased HKT1 expression (**Fig. 4B**), suggesting a positive feedback loop between TMAC2 and HKT1. Additionally, co-expression of TMAC2 and HKT1 resulted in increased levels of ABI4 and ABI5 transcription factors (**Fig. 4C**). Although ABI4 and ABI5 were not induced in the presence of TMAC2 alone, these results suggest that co-expression of TMAC2 and HKT1 beyond a specific threshold results in a feed-forward loop that further enhances HKT1 expression through activation of ABI transcription factors. Disruption of *ABI4* was previously shown to increase salt tolerance whilst its overexpression promoted salt sensitivity, and ABI4 was observed to bind to HKT1 promoter *in-vivo* (Shkolnik-Inbar, Adler, and Bar-Zvi 2013). Interestingly, we have observed an opposite effect of TMAC2 in the C24 background, where TMAC2 overexpression reduced ABA accumulation in lines with HKT1 overexpression (**Fig. 3D**). These results point to alternative signaling responses in C24 and Col-0 backgrounds, originating either from the differences in HKT1 promoter (Baxter et al. 2010; M. M. Julkowska et al. 2017), other regulatory cis-elements, or molecular elements facilitating the above described feedback between TMAC2 and HKT1 (Shkolnik-Inbar, Adler, and Bar-Zvi 2013). Future studies will be needed to reveal the specific nature of this interaction. Our results suggest that while TMAC2 is indeed involved in ABA regulation, directionality of its regulation is highly dependent on molecular context.

Tonoplast intrinsic proteins (TIPs) contribute to water and neutral solute transport (Li Song et al. 2016; Boursiac et al. 2005; Maurel et al. 1995; Pou et al. 2013), and subsets of TIPs might also contribute to ion transport (Hoai et al. 2023; Tyerman et al. 2021). TIP1;1, TIP1;2 and TIP2;2 were previously associated with facilitating lateral root emergence (Reinhardt et al. 2016), however they were thus far not identified to play a role under salt stress. Our study identified TIP2;2 to be upregulated in response to both salt treatments and HKT1 upregulation (**Fig. 2M**), with the loss-of-function TIP2;2 mutant showing increased lateral root development, and overexpression TIP2;2 lines displaying repression of lateral root development (**Fig. 5**). We have observed TIP2;2 to function as a candidate for conferring permeability to a variety of solutes and ions, and identified TIP2;2 to have close functional resemblance to its homolog TIP2;1 (**Fig. 7**). Previous studies identified tomato *SlTIP2;2*, which is most similar to *AtTIP2;2*, to regulate whole-plant transpiration, relative water content and at the cellular level, to increase the osmotic water permeability (Sade et al. 2009). While we did not study the transpiration of generated TIP2;2 mutants, it is generally known that transpiration and root development are tightly linked on a developmental level (Hepworth et al. 2016; Chater et al. 2014; Lai and Katul 2000; Sakurai-Ishikawa et al. 2011). The increase in lateral root length would increase a plant’s capability to scavenge for water, nutrients and salt, and would result in altered transpiration rates (Hepworth et al. 2016; Johannes Daniel Scharwies 2017). The subcellular localization of TIP2;2 in both plasma and tonoplast-membranes (**Fig. 6**) can provide additional level of protection under salt stress, ensuring not only high retention of sodium ions in the roots, but also their compartmentalization into the vacuole (Salazar et al. 2024; Mcgaughey et al. 2018). As TIP2;2 expression was not observed to be notable in lateral root primordia (**Fig. 6**), it would not be expected to directly affect the developing meristems and new lateral root development in a similar fashion to HKT1. Further studies investigating the contribution of TIP2;2 to HKT1-derived salt tolerance across a wider developmental context will reveal how tonoplast-intrinsic proteins can help further boost salinity resilience.

In conclusion, our study provides new insights into the complex regulatory networks involved in root development under salt stress, particularly the antagonistic interaction between HKT1 and TMAC2 and their impact on ABA signaling and lateral root outgrowth. The differential responses observed in Col-0 and C24 backgrounds underscore the importance of genetic context and tissue-specific expression in shaping transcriptional responses to stress. Additionally, our findings highlight the potential role of TIP2;2 in modulating salt tolerance through solute transport and vacuolar compartmentalization, offering a novel perspective on its function beyond water transport. These results not only advance our understanding of salt stress responses in plants but also lay the groundwork for future studies aimed at unraveling the contributions of these genes and proteins across broader developmental contexts and genetic backgrounds.

## Acknowledgments

Funding for N.A., V.J.M, J.Y.W, S.A., M.A.T, M.M.J was provided through KAUST baseline funding for M.A.T, and BTI startup funding for M.M.J.; C.S.B and A.M. were supported by the Australian Research Council (FT180100476). Additionally we would like to thank Dr. Chiou and Dr. Schaffner for sharing with us their genetic materials on PHT5:1 and UGT76B1 respectively.

## References

Ali, Akhtar, Veselin Petrov, Dae-Jin Yun, and Tsanko Gechev. 2023. “Revisiting Plant Salt Tolerance: Novel Components of the SOS Pathway.” Trends in Plant Science 28 (9): 1060– 69.

Arif, Mohammad Rashid, M. Thoihidul Islam, and Arif Hasan Khan Robin. 2019. “Salinity Stress Alters Root Morphology and Root Hair Traits in Brassica Napus.” Plants 8 (7). 10.3390/plants8070192.

Awlia, Mariam, Nouf Alshareef, Noha Saber, Arthur Korte, Helena Oakey, Klára Panzarová, Martin Trtílek, Sónia Negrão, Mark Tester, and Magdalena M. Julkowska. 2021. “Genetic Mapping of the Early Responses to Salt Stress in Arabidopsis Thaliana.” The Plant Journal: For Cell and Molecular Biology 107 (2): 544–63.

Bao, Yun, Pooja Aggarwal, Neil E. Robbins 2nd, Craig J. Sturrock, Mark C. Thompson, Han Qi Tan, Cliff Tham, et al. 2014. “Plant Roots Use a Patterning Mechanism to Position Lateral Root Branches toward Available Water.” Proceedings of the National Academy of Sciences of the United States of America 111 (25): 9319–24.

Barah, Pankaj, Mahantesha Naika B N, Naresh Doni Jayavelu, Ramanathan Sowdhamini, Khader Shameer, and Atle M. Bones. 2016. “Transcriptional Regulatory Networks in Arabidopsis Thaliana during Single and Combined Stresses.” Nucleic Acids Research 44 (7): 3147–64.

Baxter, Ivan, Jessica N. Brazelton, Danni Yu, Yu S. Huang, Brett Lahner, Elena Yakubova, Yan Li, et al. 2010. “A Coastal Cline in Sodium Accumulation in Arabidopsis Thaliana Is Driven by Natural Variation of the Sodium Transporter AtHKT1;1.” PLoS Genetics 6 (11): e1001193.

Bienert, Gerd P., Anders L. B. Møller, Kim A. Kristiansen, Alexander Schulz, Ian M. Møller, Jan K. Schjoerring, and Thomas P. Jahn. 2007. “Specific Aquaporins Facilitate the Diffusion of Hydrogen Peroxide across Membranes.” The Journal of Biological Chemistry 282 (2): 1183–92.

Bjornson, Marta, Abhaya Dandekar, and Katayoon Dehesh. 2016. “Determinants of Timing and Amplitude in the Plant General Stress Response.” Journal of Integrative Plant Biology 58 (2): 119–26.

Boursiac, Yann, Sheng Chen, Doan-Trung Luu, Mathias Sorieul, Niels van den Dries, and Christophe Maurel. 2005. “Early Effects of Salinity on Water Transport in Arabidopsis Roots. Molecular and Cellular Features of Aquaporin Expression.” Plant Physiology 139 (2): 790–805.

Byrt, Caitlin S., J. Damien Platten, Wolfgang Spielmeyer, Richard A. James, Evans S. Lagudah, Elizabeth S. Dennis, Mark Tester, and Rana Munns. 2007. “HKT1;5-like Cation Transporters Linked to Na+ Exclusion Loci in Wheat, Nax2 and Kna1.” Plant Physiology 143 (4): 1918–28.

Casimiro, Ilda, Tom Beeckman, Neil Graham, Rishikesh Bhalerao, Hanma Zhang, Pedro Casero, Goran Sandberg, and Malcolm J. Bennett. 2003. “Dissecting Arabidopsis Lateral Root Development.” Trends in Plant Science 8 (4): 165–71.

Chater, Caspar C. C., James Oliver, Stuart Casson, and Julie E. Gray. 2014. “Putting the Brakes on: Abscisic Acid as a Central Environmental Regulator of Stomatal Development.” The New Phytologist 202 (2): 376–91.

Choi, Won-Gyu, Masatsugu Toyota, Su-Hwa Kim, Richard Hilleary, and Simon Gilroy. 2014. “Salt Stress-Induced Ca2+ Waves Are Associated with Rapid, Long-Distance Root-to-Shoot Signaling in Plants.” Proceedings of the National Academy of Sciences of the United States of America 111 (17): 6497–6502.

Cuin, Tracey Ann, and Sergey Shabala. 2007. “Amino Acids Regulate Salinity-Induced Potassium Efflux in Barley Root Epidermis.” Planta 225 (3): 753–61.

Das, Sanjukta, and Manju Bansal. 2019. “Variation of Gene Expression in Plants Is Influenced by Gene Architecture and Structural Properties of Promoters.” PloS One 14 (3): e0212678.

Davenport, Romola Jane, Alicia Muñoz-Mayor, Deepa Jha, Pauline Adobea Essah, Ana Rus, and Mark Tester. 2007. “The Na+ Transporter AtHKT1;1 Controls Retrieval of Na+ from the Xylem in Arabidopsis.” Plant, Cell & Environment 30 (4): 497–507.

De Rybel, Bert, Valya Vassileva, Boris Parizot, Marlies Demeulenaere, Wim Grunewald, Dominique Audenaert, Jelle Van Campenhout, et al. 2010. “A Novel aux/IAA28 Signaling Cascade Activates GATA23-Dependent Specification of Lateral Root Founder Cell Identity.” Current Biology: CB 20 (19): 1697–1706.

De Smet, Ive, Laurent Signora, Tom Beeckman, Dirk Inzé, Christine H. Foyer, and Hanma Zhang. 2003. “An Abscisic Acid-Sensitive Checkpoint in Lateral Root Development of Arabidopsis.” The Plant Journal: For Cell and Molecular Biology 33 (3): 543–55.

De Smet, Ive, Takuya Tetsumura, Bert De Rybel, Nicolas Frei dit Frey, Laurent Laplaze, Ilda Casimiro, Ranjan Swarup, et al. 2007. “Auxin-Dependent Regulation of Lateral Root Positioning in the Basal Meristem of Arabidopsis.” Development 134 (4): 681–90.

Duan, Lina, Daniela Dietrich, Chong Han Ng, Penny Mei Yeen Chan, Rishikesh Bhalerao, Malcolm J. Bennett, and José R. Dinneny. 2013. “Endodermal ABA Signaling Promotes Lateral Root Quiescence during Salt Stress in Arabidopsis Seedlings.” The Plant Cell 25 (1): 324–41.

Dubrovsky, J. G., P. W. Doerner, A. Colón-Carmona, and T. L. Rost. 2000. “Pericycle Cell Proliferation and Lateral Root Initiation in Arabidopsis.” Plant Physiology 124 (4): 1648–57.

Evans, Matthew J., Won-Gyu Choi, Simon Gilroy, and Richard J. Morris. 2016. “A ROS-Assisted Calcium Wave Dependent on the AtRBOHD NADPH Oxidase and TPC1 Cation Channel Propagates the Systemic Response to Salt Stress.” Plant Physiology 171 (3): 1771–84.

Feng, Wei, Daniel Kita, Alexis Peaucelle, Heather N. Cartwright, Vinh Doan, Qiaohong Duan, Ming-Che Liu, et al. 2018. “The FERONIA Receptor Kinase Maintains Cell-Wall Integrity during Salt Stress through Ca2+ Signaling.” Current Biology: CB 28 (5): 666–75.e5.

Finkina, E. I., D. N. Melnikova, I. V. Bogdanov, and T. V. Ovchinnikova. 2016. “Lipid Transfer Proteins As Components of the Plant Innate Immune System: Structure, Functions, and Applications.” Acta Naturae 8 (2): 47–61.

Garcia, Mary Emily, Tim Lynch, Julian Peeters, Chris Snowden, and Ruth Finkelstein. 2008. “A Small Plant-Specific Protein Family of ABI Five Binding Proteins (AFPs) Regulates Stress Response in Germinating Arabidopsis Seeds and Seedlings.” Plant Molecular Biology 67 (6): 643–58.

Geldner, Niko, Valérie Dénervaud-Tendon, Derek L. Hyman, Ulrike Mayer, York-Dieter Stierhof, and Joanne Chory. 2009. “Rapid, Combinatorial Analysis of Membrane Compartments in Intact Plants with a Multicolor Marker Set.” The Plant Journal: For Cell and Molecular Biology 59 (1): 169–78.

Goh, Tatsuaki, Shunpei Joi, Tetsuro Mimura, and Hidehiro Fukaki. 2012. “The Establishment of Asymmetry in Arabidopsis Lateral Root Founder Cells Is Regulated by LBD16/ASL18 and Related LBD/ASL Proteins.” Development 139 (5): 883–93.

Groszmann, Michael, Annamaria De Rosa, Weihua Chen, Jiaen Qiu, Samantha A. McGaughey, Caitlin S. Byrt, and John R. Evans. 2023. “A High-Throughput Yeast Approach to Characterize Aquaporin Permeabilities: Profiling the Arabidopsis PIP Aquaporin Sub-Family.” Frontiers in Plant Science 14 (January):1078220.

Han, Xiao, Chenxu Gao, Lisen Liu, Yihao Zhang, Yuying Jin, Qingdi Yan, Lan Yang, Fuguang Li, and Zhaoen Yang. 2022. “Integration of eQTL Analysis and GWAS Highlights Regulation Networks in Cotton under Stress Condition.” International Journal of Molecular Sciences 23 (14). 10.3390/ijms23147564.

Hepworth, Christopher, Carla Turner, Marcela Guimaraes Landim, Duncan Cameron, and Julie E. Gray. 2016. “Balancing Water Uptake and Loss through the Coordinated Regulation of Stomatal and Root Development.” PloS One 11 (6): e0156930.

Hoai, Phan Thi Thanh, Jiaen Qiu, Michael Groszmann, Annamaria De Rosa, Stephen D. Tyerman, and Caitlin S. Byrt. 2023. “Arabidopsis Plasma Membrane Intrinsic Protein (AtPIP2;1) Is Implicated in a Salinity Conditional Influence on Seed Germination.” Functional Plant Biology: FPB 50 (8): 633–48.

Huang, Ming-Der, and Wen-Luan Wu. 2007. “Overexpression of TMAC2, a Novel Negative Regulator of Abscisic Acid and Salinity Responses, Has Pleiotropic Effects in Arabidopsis Thaliana.” Plant Molecular Biology 63 (4): 557–69.

Ibeas, Miguel Angel, Hernán Salinas-Grenet, Nathan R. Johnson, Jorge Pérez-Díaz, Elena A. Vidal, José Miguel Alvarez, and José M. Estevez. 2024. “Filling the Gaps on Root Hair Development under Salt Stress and Phosphate Starvation Using Current Evidence Coupled with a Meta-Analysis Approach.” Plant Physiology, June. 10.1093/plphys/kiae346.

Ishka, Maryam Rahmati, Hayley Sussman, Yunfei Hu, Mashael Daghash Alqahtani, Eric Craft, Ronell Sicat, Minmin Wang, et al. 2024. “Natural Variation in Salt-Induced Changes in Root:shoot Ratio Reveals SR3G as a Negative Regulator of Root Suberization and Salt Resilience in Arabidopsis.” bioRxiv. 10.1101/2024.04.09.588564.

Jaime-Pérez, Noelia, Benito Pineda, Begoña García-Sogo, Alejandro Atares, Asmini Athman, Caitlin S. Byrt, Raquel Olías, et al. 2017. “The Sodium Transporter Encoded by the HKT1;2 Gene Modulates Sodium/potassium Homeostasis in Tomato Shoots under Salinity.” Plant, Cell & Environment 40 (5): 658–71.

Julkowska, M. 2020. “mRNA Purification with Ambion Dynabeads,” January. 10.17504/protocols.io.pbrdim6.

Julkowska, Magdalena. 2018. “RNA Isolation with TRIzol v1.” Protocols.io. ZappyLab, Inc. 10.17504/protocols.io.pbndime.

Julkowska, Magdalena M., Huub C. J. Hoefsloot, Selena Mol, Richard Feron, Gert-Jan de Boer, Michel A. Haring, and Christa Testerink. 2014. “Capturing Arabidopsis Root Architecture Dynamics with ROOT-FIT Reveals Diversity in Responses to Salinity.” Plant Physiology 166 (3): 1387–1402.

Julkowska, Magdalena M., Iko T. Koevoets, Selena Mol, Huub Hoefsloot, Richard Feron, Mark A. Tester, Joost J. B. Keurentjes, et al. 2017. “Genetic Components of Root Architecture Remodeling in Response to Salt Stress.” The Plant Cell 29 (12): 3198–3213.

Kawa, Dorota, Magdalena M. Julkowska, Hector Montero Sommerfeld, Anneliek Ter Horst, Michel A. Haring, and Christa Testerink. 2016. “Phosphate-Dependent Root System Architecture Responses to Salt Stress.” Plant Physiology 172 (2): 690–706.

Koncz, Csaba, and Jeff Schell. 1986. “The Promoter of TL-DNA Gene 5 Controls the Tissue-Specific Expression of Chimaeric Genes Carried by a Novel Type of Agrobacterium Binary Vector.” Molecular & General Genetics: MGG 204 (3): 383–96.

Lai, Chun-Ta, and Gabriel Katul. 2000. “The Dynamic Role of Root-Water Uptake in Coupling Potential to Actual Transpiration.” Advances in Water Resources 23 (4): 427–39.

Lampropoulos, Athanasios, Zoran Sutikovic, Christian Wenzl, Ira Maegele, Jan U. Lohmann, and Joachim Forner. 2013. “GreenGate---a Novel, Versatile, and Efficient Cloning System for Plant Transgenesis.” PloS One 8 (12): e83043.

Lee, J., C. Godon, G. Lagniel, D. Spector, J. Garin, J. Labarre, and M. B. Toledano. 1999. “Yap1 and Skn7 Control Two Specialized Oxidative Stress Response Regulons in Yeast.” The Journal of Biological Chemistry 274 (23): 16040–46.

Lin, Meng, Pengfei Qiao, Susanne Matschi, Miguel Vasquez, Guillaume P. Ramstein, Richard Bourgault, Marc Mohammadi, et al. 2022. “Integrating GWAS and TWAS to Elucidate the Genetic Architecture of Maize Leaf Cuticular Conductance.” Plant Physiology 189 (4): 2144–58.

Liu, Lai-Hua, Uwe Ludewig, Brigitte Gassert, Wolf B. Frommer, and Nicolaus von Wirén. 2003. “Urea Transport by Nitrogen-Regulated Tonoplast Intrinsic Proteins in Arabidopsis.” Plant Physiology 133 (3): 1220–28.

Liu, Tzu-Yin, Teng-Kuei Huang, Shu-Yi Yang, Yu-Ting Hong, Sheng-Min Huang, Fu-Nien Wang, Su-Fen Chiang, Shang-Yueh Tsai, Wen-Chien Lu, and Tzyy-Jen Chiou. 2016. “Identification of Plant Vacuolar Transporters Mediating Phosphate Storage.” Nature Communications 7 (1): 11095.

Lobet, Guillaume, Loïc Pagès, and Xavier Draye. 2011. “A Novel Image-Analysis Toolbox Enabling Quantitative Analysis of Root System Architecture.” Plant Physiology 157 (1): 29– 39.

Loqué, Dominique, Uwe Ludewig, Lixing Yuan, and Nicolaus von Wirén. 2005. “Tonoplast Intrinsic Proteins AtTIP2;1 and AtTIP2;3 Facilitate NH3 Transport into the Vacuole.” Plant Physiology 137 (2): 671–80.

Lynch, Tim J., B. Joy Erickson, Dusty R. Miller, and Ruth R. Finkelstein. 2017. “ABI5-Binding Proteins (AFPs) Alter Transcription of ABA-Induced Genes via a Variety of Interactions with Chromatin Modifiers.” Plant Molecular Biology 93 (4-5): 403–18.

Mäser, Pascal, Yoshihiro Hosoo, Shinobu Goshima, Tomoaki Horie, Brendan Eckelman, Katsuyuki Yamada, Kazuya Yoshida, et al. 2002. “Glycine Residues in Potassium Channel-like Selectivity Filters Determine Potassium Selectivity in Four-Loop-per-Subunit HKT Transporters from Plants.” Proceedings of the National Academy of Sciences of the United States of America 99 (9): 6428–33.

Maurel, C., M. Chrispeels, C. Lurin, F. Tacnet, D. Geelen, P. Ripoche, and J. Guern. 1997. “Function and Regulation of Seed Aquaporins.” Journal of Experimental Botany 48 Spec No (Special): 421–30.

Maurel, C., R. T. Kado, J. Guern, and M. J. Chrispeels. 1995. “Phosphorylation Regulates the Water Channel Activity of the Seed-Specific Aquaporin Alpha-TIP.” The EMBO Journal 14 (13): 3028–35.

Ma, Yizan, Ling Min, Junduo Wang, Yaoyao Li, Yuanlong Wu, Qin Hu, Yuanhao Ding, et al. 2021. “A Combination of Genome-Wide and Transcriptome-Wide Association Studies Reveals Genetic Elements Leading to Male Sterility during High Temperature Stress in Cotton.” The New Phytologist 231 (1): 165–81.

Mcgaughey, S. A., J. Qiu, S. D. Tyerman, and C. S. Byrt. 2018. “Regulating Root Aquaporin Function in Response to Changes in Salinity.” Annual Plant Reviews Online 1 (2): 1–36.

Mehra, Poonam, Bipin K. Pandey, Dalia Melebari, Jason Banda, Nicola Leftley, Valentin Couvreur, James Rowe, et al. 2022. “Hydraulic Flux–responsive Hormone Redistribution Determines Root Branching.” Science 378 (6621): 762–68.

Miura, Kenji, Aiko Sato, Masaru Ohta, and Jun Furukawa. 2011. “Increased Tolerance to Salt Stress in the Phosphate-Accumulating Arabidopsis Mutants siz1 and pho2.” Planta 234 (6): 1191–99.

Møller, Inge S., Matthew Gilliham, Deepa Jha, Gwenda M. Mayo, Stuart J. Roy, Juliet C. Coates, Jim Haseloff, and Mark Tester. 2009. “Shoot Na+ Exclusion and Increased Salinity Tolerance Engineered by Cell Type-Specific Alteration of Na+ Transport in Arabidopsis.” The Plant Cell 21 (7): 2163–78.

Munns, Rana, Richard A. James, Bo Xu, Asmini Athman, Simon J. Conn, Charlotte Jordans, Caitlin S. Byrt, et al. 2012. “Wheat Grain Yield on Saline Soils Is Improved by an Ancestral Na+ Transporter Gene.” Nature Biotechnology 30 (4): 360–64.

Orman-Ligeza, Beata, Emily C. Morris, Boris Parizot, Tristan Lavigne, Aurelie Babé, Aleksander Ligeza, Stephanie Klein, et al. 2018. “The Xerobranching Response Represses Lateral Root Formation When Roots Are Not in Contact with Water.” Current Biology: CB 28 (19): 3165–73.e5.

Orosa-Puente, Beatriz, Nicola Leftley, Daniel von Wangenheim, Jason Banda, Anjil K. Srivastava, Kristine Hill, Jekaterina Truskina, et al. 2018. “Root Branching toward Water Involves Posttranslational Modification of Transcription Factor ARF7.” Science 362 (6421): 1407–10.

Parizot, Boris, Laurent Laplaze, Lilian Ricaud, Elodie Boucheron-Dubuisson, Vincent Bayle, Martin Bonke, Ive De Smet, et al. 2008. “Diarch Symmetry of the Vascular Bundle in Arabidopsis Root Encompasses the Pericycle and Is Reflected in Distich Lateral Root Initiation.” Plant Physiology 146 (1): 140–48.

Pignon, Charles P., Samuel B. Fernandes, Ravi Valluru, Nonoy Bandillo, Roberto Lozano, Edward Buckler, Michael A. Gore, Stephen P. Long, Patrick J. Brown, and Andrew D. B. Leakey. 2021. “Phenotyping Stomatal Closure by Thermal Imaging for GWAS and TWAS of Water Use Efficiency-Related Genes.” Plant Physiology 187 (4): 2544–62.

Pou, Alicia, Hipolito Medrano, Jaume Flexas, and Stephen D. Tyerman. 2013. “A Putative Role for TIP and PIP Aquaporins in Dynamics of Leaf Hydraulic and Stomatal Conductances in Grapevine under Water Stress and Re-Watering.” Plant, Cell & Environment 36 (4): 828– 43.

Quintero, F. J., M. R. Blatt, and J. M. Pardo. 2000. “Functional Conservation between Yeast and Plant Endosomal Na(+)/H(+) Antiporters.” FEBS Letters 471 (2-3): 224–28.

Ramakrishna, Priya, Paola Ruiz Duarte, Graham A. Rance, Martin Schubert, Vera Vordermaier, Lam Dai Vu, Evan Murphy, et al. 2019. “EXPANSIN A1-Mediated Radial Swelling of Pericycle Cells Positions Anticlinal Cell Divisions during Lateral Root Initiation.” Proceedings of the National Academy of Sciences of the United States of America 116 (17): 8597–8602.

Reinhardt, Hagen, Charles Hachez, Manuela Désirée Bienert, Azeez Beebo, Kamal Swarup, Ute Voß, Karim Bouhidel, et al. 2016. “Tonoplast Aquaporins Facilitate Lateral Root Emergence.” Plant Physiology 170 (3): 1640–54.

Rus, Ana, Ivan Baxter, Balasubramaniam Muthukumar, Jeff Gustin, Brett Lahner, Elena Yakubova, and David E. Salt. 2006. “Natural Variants of AtHKT1 Enhance Na+ Accumulation in Two Wild Populations of Arabidopsis.” PLoS Genetics 2 (12): e210.

Sade, Nir, Basia J. Vinocur, Alex Diber, Arava Shatil, Gil Ronen, Hagit Nissan, Rony Wallach, Hagai Karchi, and Menachem Moshelion. 2009. “Improving Plant Stress Tolerance and Yield Production: Is the Tonoplast Aquaporin SlTIP2;2 a Key to Isohydric to Anisohydric Conversion?” The New Phytologist 181 (3): 651–61.

Saint Paul, Veronica von, Wei Zhang, Basem Kanawati, Birgit Geist, Theresa Faus-Kessler, Philippe Schmitt-Kopplin, and Anton R. Schäffner. 2011. “The Arabidopsis Glucosyltransferase UGT76B1 Conjugates Isoleucic Acid and Modulates Plant Defense and Senescence.” The Plant Cell 23 (11): 4124–45.

Sakurai-Ishikawa, Junko, Mari Murai-Hatano, Hidehiro Hayashi, Arifa Ahamed, Keiko Fukushi, Tadashi Matsumoto, and Yoshichika Kitagawa. 2011. “Transpiration from Shoots Triggers Diurnal Changes in Root Aquaporin Expression.” Plant, Cell & Environment 34 (7): 1150– 63.

Salazar, Octavio R., Ke Chen, Vanessa J. Melino, Muppala P. Reddy, Eva Hribová, Jana Cížková, Denisa Beránková, et al. 2024. “SOS1 Tonoplast Neo-Localization and the RGG Protein SALTY Are Important in the Extreme Salinity Tolerance of Salicornia Bigelovii.” Nature Communications 15 (1): 4279.

Scharwies, Johannes Daniel. 2017. “The Role of Aquaporins in Plant Responses to Drought.” http://hdl.handle.net/2440/113120.

Scharwies, Johannes D., Taylor Clarke, Zihao Zheng, Andrea Dinneny, Siri Birkeland, Margaretha A. Veltman, Craig J. Sturrock, et al. 2024. “Maize Genetic Diversity Identifies Moisture-Dependent Root-Branch Signaling Pathways.” Plant Biology. bioRxiv. https://www.biorxiv.org/content/10.1101/2024.08.26.609741v1.full.

Shamaya, Nawar Jalal, Yuri Shavrukov, Peter Langridge, Stuart John Roy, and Mark Tester. “Genetics of Na+ Exclusion and Salinity Tolerance in Afghani Durum Wheat Landraces.” BMC Plant Biology 17 (1): 209.

Shkolnik-Inbar, Doron, Guy Adler, and Dudy Bar-Zvi. 2013. “ABI4 Downregulates Expression of the Sodium Transporter HKT1;1 in Arabidopsis Roots and Affects Salt Tolerance.” The Plant Journal: For Cell and Molecular Biology 73 (6): 993–1005.

Simon, Aaron D., and Susan Gitler. 2007. “A Suite of Gateway® Cloning Vectors for High-Throughput Genetic Analysis in Saccharomyces Cerevisiae.” Yeast 24 (10): 913–19.

Song, Huifang, Yibo Cao, Xinyan Zhao, and Lingyun Zhang. 2023. “Na+-Preferential Ion Transporter HKT1;1 Mediates Salt Tolerance in Blueberry.” Plant Physiology 194 (1): 511– 29.

Song, Liang, Shao-Shan Carol Huang, Aaron Wise, Rosa Castanon, Joseph R. Nery, Huaming Chen, Marina Watanabe, Jerushah Thomas, Ziv Bar-Joseph, and Joseph R. Ecker. 2016. “A Transcription Factor Hierarchy Defines an Environmental Stress Response Network.” Science 354 (6312). 10.1126/science.aag1550.

Song, Li, Na Nguyen, Rupesh K. Deshmukh, Gunvant B. Patil, Silvas J. Prince, Babu Valliyodan, Raymond Mutava, Sharon M. Pike, Walter Gassmann, and Henry T. Nguyen. 2016. “Soybean TIP Gene Family Analysis and Characterization of GmTIP1;5 and GmTIP2;5 Water Transport Activity.” Frontiers in Plant Science 7 (October):1564.

Song, Yushuang, Simin Li, Yi Sui, Hongxiang Zheng, Guoliang Han, Xi Sun, Wenjing Yang, et al. 2022. “SbbHLH85, a bHLH Member, Modulates Resilience to Salt Stress by Regulating Root Hair Growth in Sorghum.” TAG. Theoretical and Applied Genetics. Theoretische Und Angewandte Genetik 135 (1): 201–16.

Sparkes, Imogen A., John Runions, Anne Kearns, and Chris Hawes. 2006. “Rapid, Transient Expression of Fluorescent Fusion Proteins in Tobacco Plants and Generation of Stably Transformed Plants.” Nature Protocols 1 (4): 2019–25.

Sudhakaran, Sreeja, Vandana Thakral, Gunashri Padalkar, Nitika Rajora, Pallavi Dhiman, Gaurav Raturi, Yogesh Sharma, et al. 2021. “Significance of Solute Specificity, Expression, and Gating Mechanism of Tonoplast Intrinsic Protein during Development and Stress Response in Plants.” Physiologia Plantarum 172 (1): 258–74.

Tanghe, An, Patrick Van Dijck, Françoise Dumortier, Aloys Teunissen, Stefan Hohmann, and Johan M. Thevelein. 2002. “Aquaporin Expression Correlates with Freeze Tolerance in Baker’s Yeast, and Overexpression Improves Freeze Tolerance in Industrial Strains.” Applied and Environmental Microbiology 68 (12): 5981–89.

Tyerman, Stephen D., Samantha A. McGaughey, Jiaen Qiu, Andrea J. Yool, and Caitlin S. Byrt. 2021. “Adaptable and Multifunctional Ion-Conducting Aquaporins.” Annual Review of Plant Biology 72 (June):703–36.

Van den Broeck, Lisa, Marieke Dubois, Mattias Vermeersch, Veronique Storme, Minami Matsui, and Dirk Inzé. 2017. “From Network to Phenotype: The Dynamic Wiring of an Arabidopsis Transcriptional Network Induced by Osmotic Stress.” Molecular Systems Biology 13 (12): 961.

Vihervaara, Anniina, Dig Bijay Mahat, Michael J. Guertin, Tinyi Chu, Charles G. Danko, John T. Lis, and Lea Sistonen. 2017. “Transcriptional Response to Stress Is Pre-Wired by Promoter and Enhancer Architecture.” Nature Communications 8 (1): 255.

Voinnet, Olivier, Susana Rivas, Pere Mestre, and David Baulcombe. 2003. “An Enhanced Transient Expression System in Plants Based on Suppression of Gene Silencing by the p19 Protein of Tomato Bushy Stunt Virus.” The Plant Journal: For Cell and Molecular Biology 33 (5): 949–56.

Wang, Jian You, Saleh Alseekh, Tingting Xiao, Abdugaffor Ablazov, Leonardo Perez de Souza, Valentina Fiorilli, Marita Anggarani, et al. 2021. “Multi-Omics Approaches Explain the Growth-Promoting Effect of the Apocarotenoid Growth Regulator Zaxinone in Rice.” Communications Biology 4 (1): 1222.

Wang, Youning, Wensheng Zhang, Kexue Li, Feifei Sun, Chunyu Han, Yukun Wang, and Xia Li. 2008. “Salt-Induced Plasticity of Root Hair Development Is Caused by Ion Disequilibrium in Arabidopsis Thaliana.” Journal of Plant Research 121 (1): 87–96.

Yang, Li, Ziqiang Liu, Feng Lu, Aiwu Dong, and Hai Huang. 2006. “SERRATE Is a Novel Nuclear Regulator in Primary microRNA Processing in Arabidopsis.” The Plant Journal: For Cell and Molecular Biology 47 (6): 841–50.

Yu, Li‘ang, Anna C. Nelson Dittrich, Xiaodan Zhang, Jordan R. Brock, Venkatesh P. Thirumalaikumar, Giovanni Melandri, Aleksandra Skirycz, et al. 2024. “Regulation of a Single Inositol 1-Phosphate Synthase Homeologue by HSFA6B Contributes to Fibre Yield Maintenance under Drought Conditions in Upland Cotton.” Plant Biotechnology Journal, June. 10.1111/pbi.14402.

Zelm, Eva van, Yanxia Zhang, and Christa Testerink. 2020. “Salt Tolerance Mechanisms of Plants.” Annual Review of Plant Biology, March. 10.1146/annurev-arplant-050718-100005.

Zou, Yutao, Nora Gigli-Bisceglia, Eva van Zelm, Pinelopi Kokkinopoulou, Magdalena M. Julkowska, Maarten Besten, Thu-Phuong Nguyen, et al. 2024. “Arabinosylation of Cell Wall Extensin Is Required for the Directional Response to Salinity in Roots.” The Plant Cell, May. 10.1093/plcell/koae135.

