## Supplemental Figures for "Root Remodeling Mechanisms and Salt Tolerance Trade-Offs: The Roles of HKT1, TMAC2, and TIP2;2 in Arabidopsis"

### Supplemental Figures and Table Legends

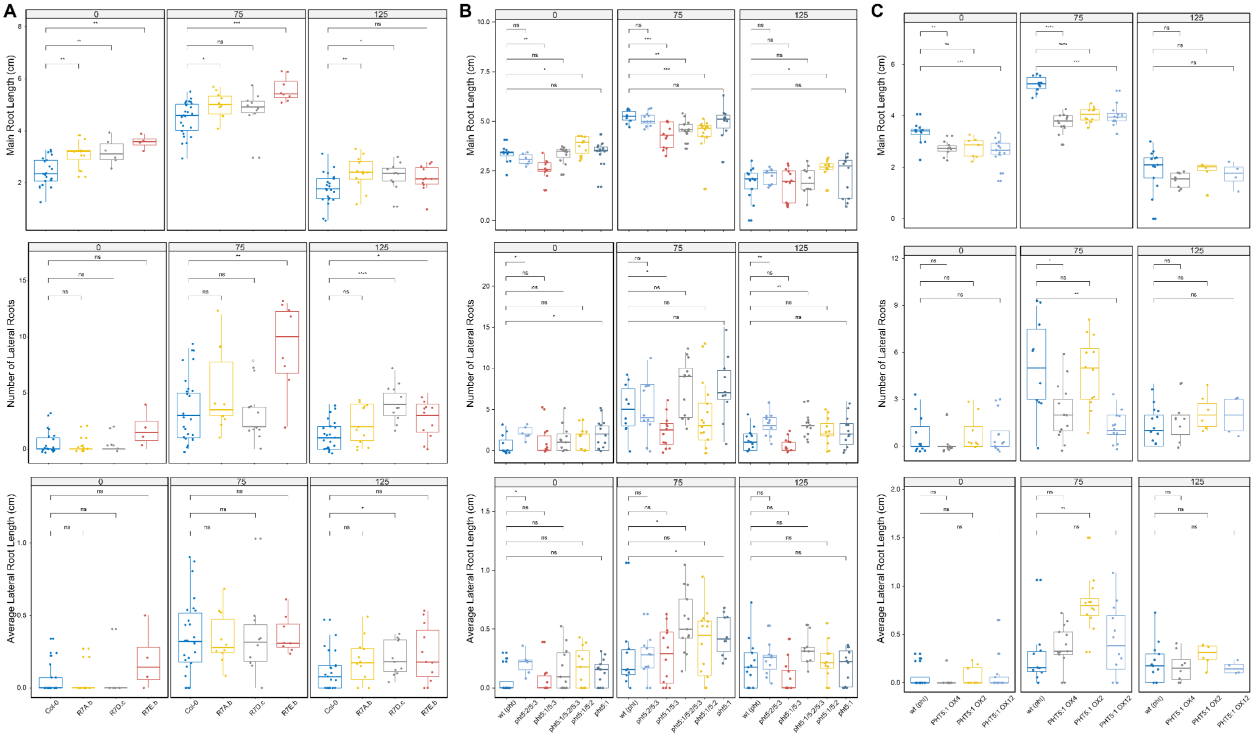


**Figure S1. Contributions of vacuolar phosphate transporter (**AT1G63010 - PHT5:1**) to salt-induced changes in root architecture.** Root System architecture of **(A)** T-DNA insertion lines (SALK_023873, SAIL_789_E03 and SALK_098188C) **(B)** Single and multiple mutant lines obtained from Dr. Chiou Lab (T.-Y. Liu et al. 2016), and **(C)** gain-of-function liens obtained from Dr. Chiou Lab (T.-Y. Liu et al. 2016). The 4 days old seedlings were exposed to control (0 mM NaCl) or salt stress treatment (75 or 125 mM NaCl) for 4 (control) and 12 days (salt stress) of treatment prior to quantification of root architecture using SmartRoot. The graphs represent individual components of root architecture: Main Root Length, lateral root number and average lateral root length. The significant differences between individual mutant lines and their respective background lines were determined using one-way ANOVA test, with *, **, *** and **** indicating p-values below 0.05, 0.01, 0.001 and 0.0001 respectively.

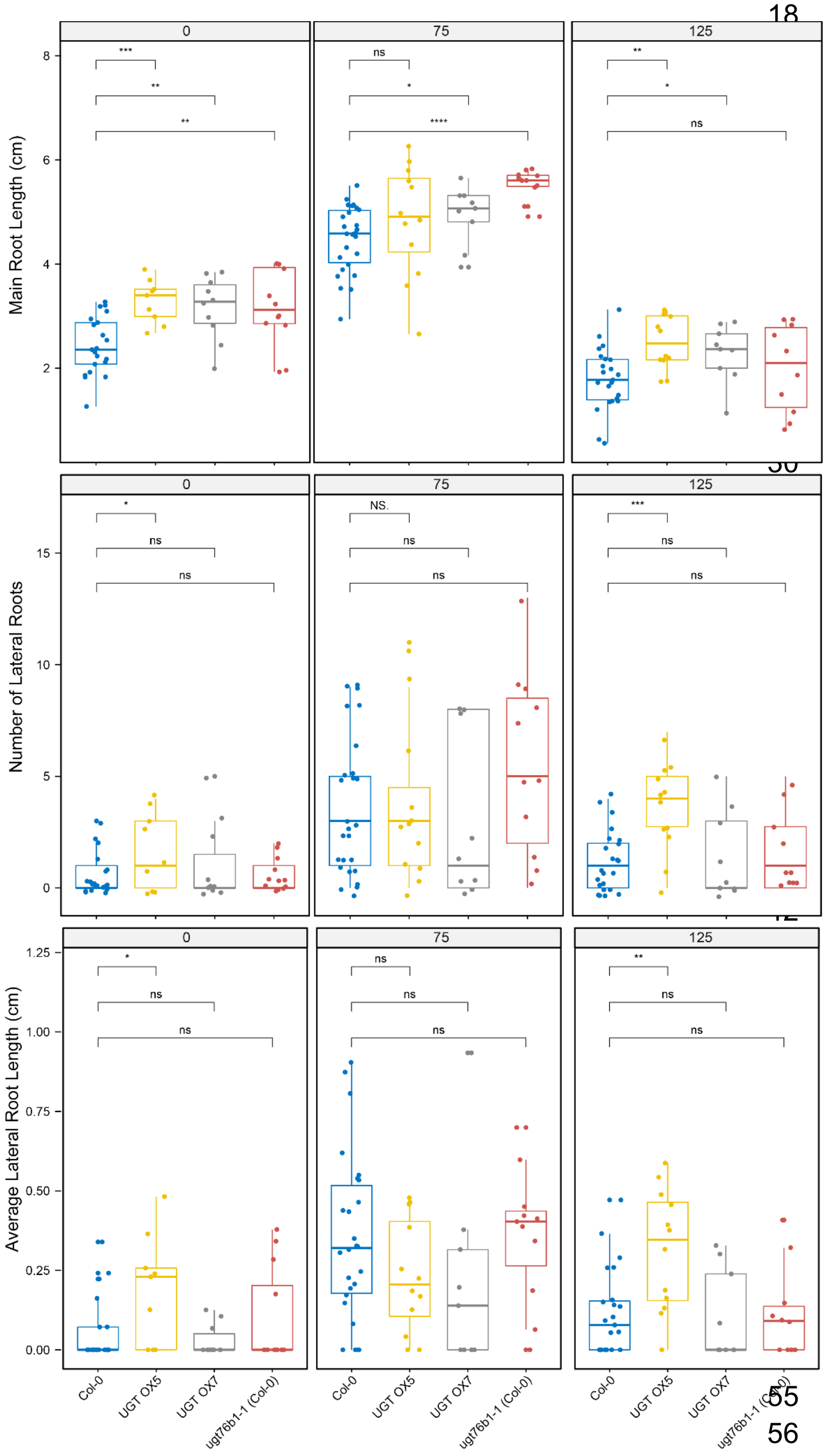


**Figure S2. Contributions of UDP-dependent glycosyltransferase 76 B1 (AT3G11340 - UGT76B1) to salt-induced changes in root architecture.** Root System architecture of two overexpression (OX) lines and one loss-of-function line obtained from Dr. Schaffner Lab (von Saint Paul et al. 2011). The 4 days old seedlings were exposed to control (0 mM NaCl) or salt stress treatment (75 or 125 mM NaCl) for 4 (control) and 12 days (salt stress) of treatment prior to quantification of root architecture using SmartRoot. The graphs represent individual components of root architecture: Main Root Length, lateral root number and average lateral root length. The significant differences between individual mutant lines and their respective background lines were determined using one-way ANOVA test, with *, **, *** and **** indicating p-values below 0.05, 0.01, 0.001 and 0.0001 respectively.

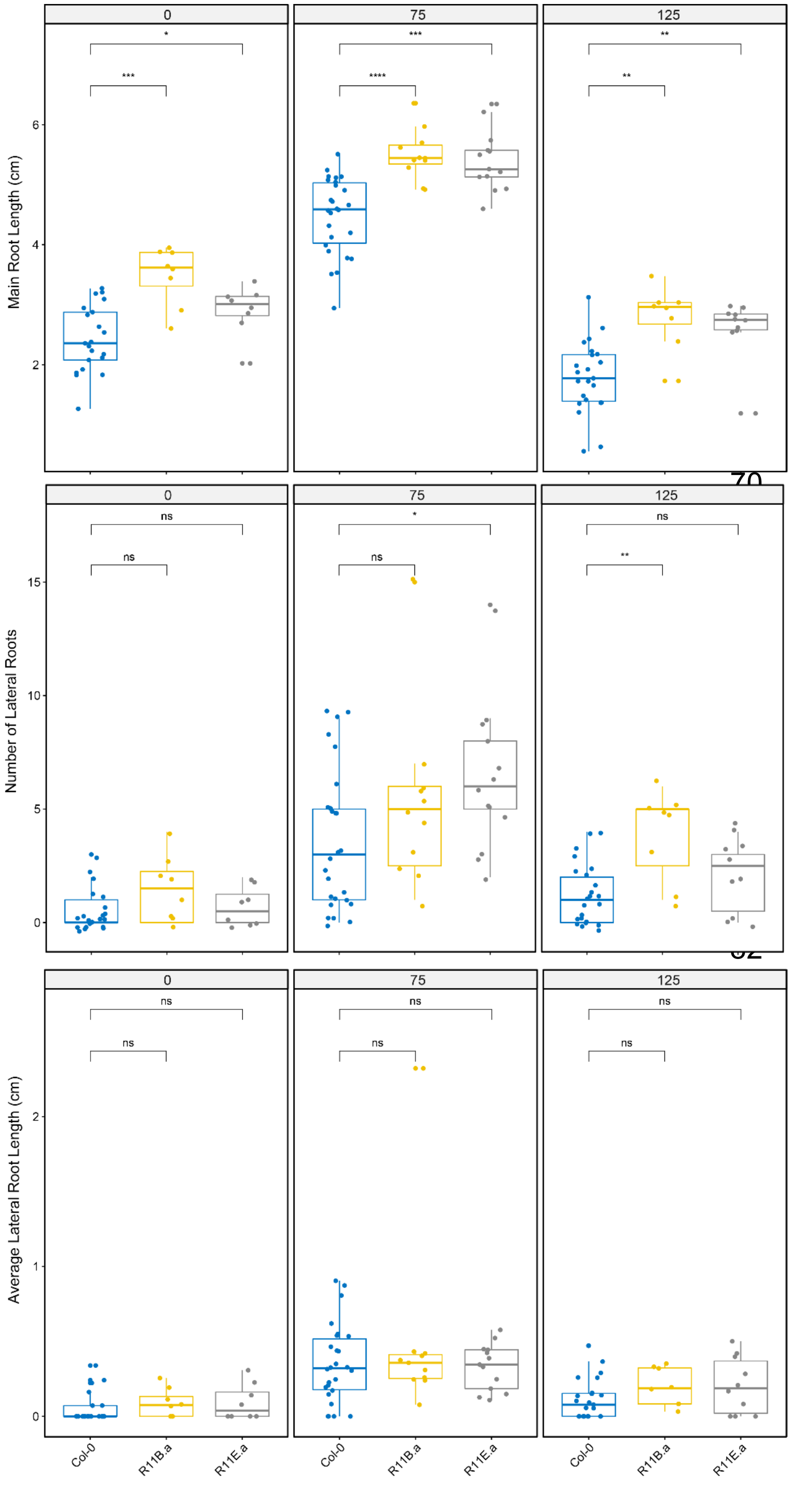


**Figure S3. Contributions of $CO_2$ responsive secreted protease (AT1G20160 - SBT5.2) to salt-induced changes in root architecture.** Root System architecture of T-DNA insertion lines (SALK_099861 and SALK_012112). The 4 days old seedlings were exposed to control (0 mM NaCl) or salt stress treatment (75 or 125 mM NaCl) for 4 (control) and 12 days (salt stress) of treatment prior to quantification of root architecture using SmartRoot. The graphs represent individual components of root architecture: Main Root Length, lateral root number and average lateral root length. The significant differences between individual mutant lines and their respective background lines were determined using one-way ANOVA test, with *, **, *** and **** indicating p-values below 0.05, 0.01, 0.001 and 0.0001 respectively.

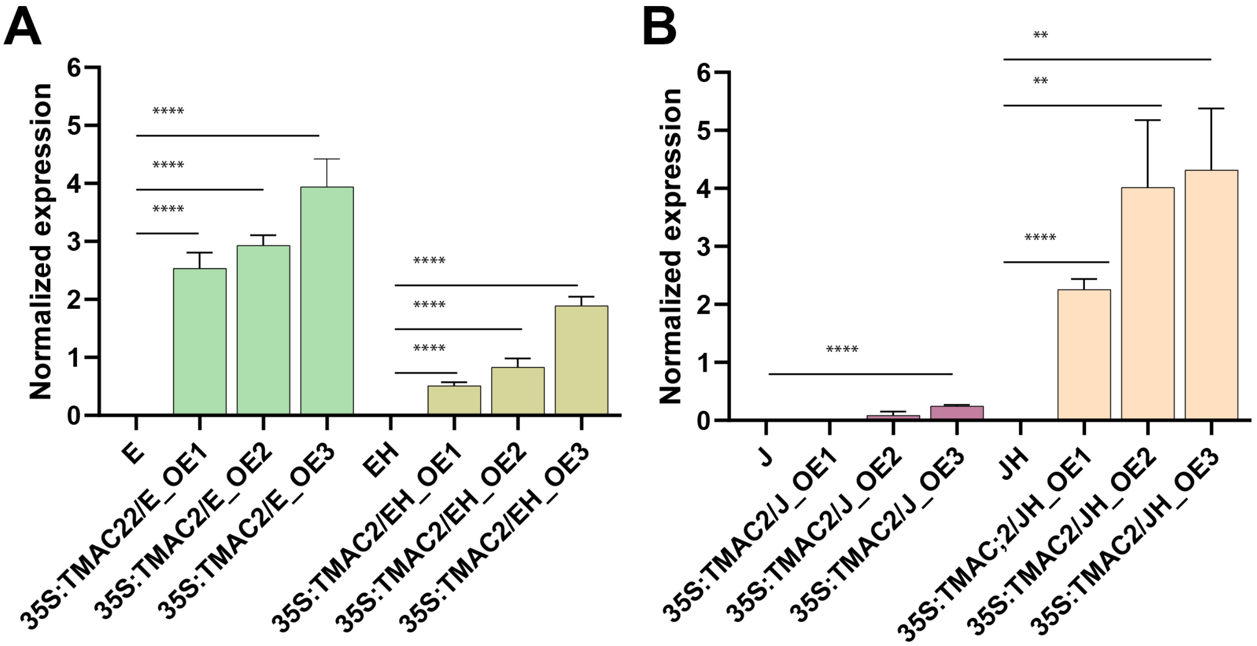


**Figure S4. Expression of TMAC2 in generated overexpression lines.** Expression of TMAC2 in Col-0 (E2586) and C24 (J2731) background with or without additional tissue-specific overexpression of HKT1. All expression results are based on three independent biological replicates collected from the leaves of soil grown plants. The significant differences between individual mutant lines and their respective background lines were determined using one-way ANOVA test, with *, **, *** and **** indicating p-values below 0.05, 0.01, 0.001 and 0.0001 respectively.

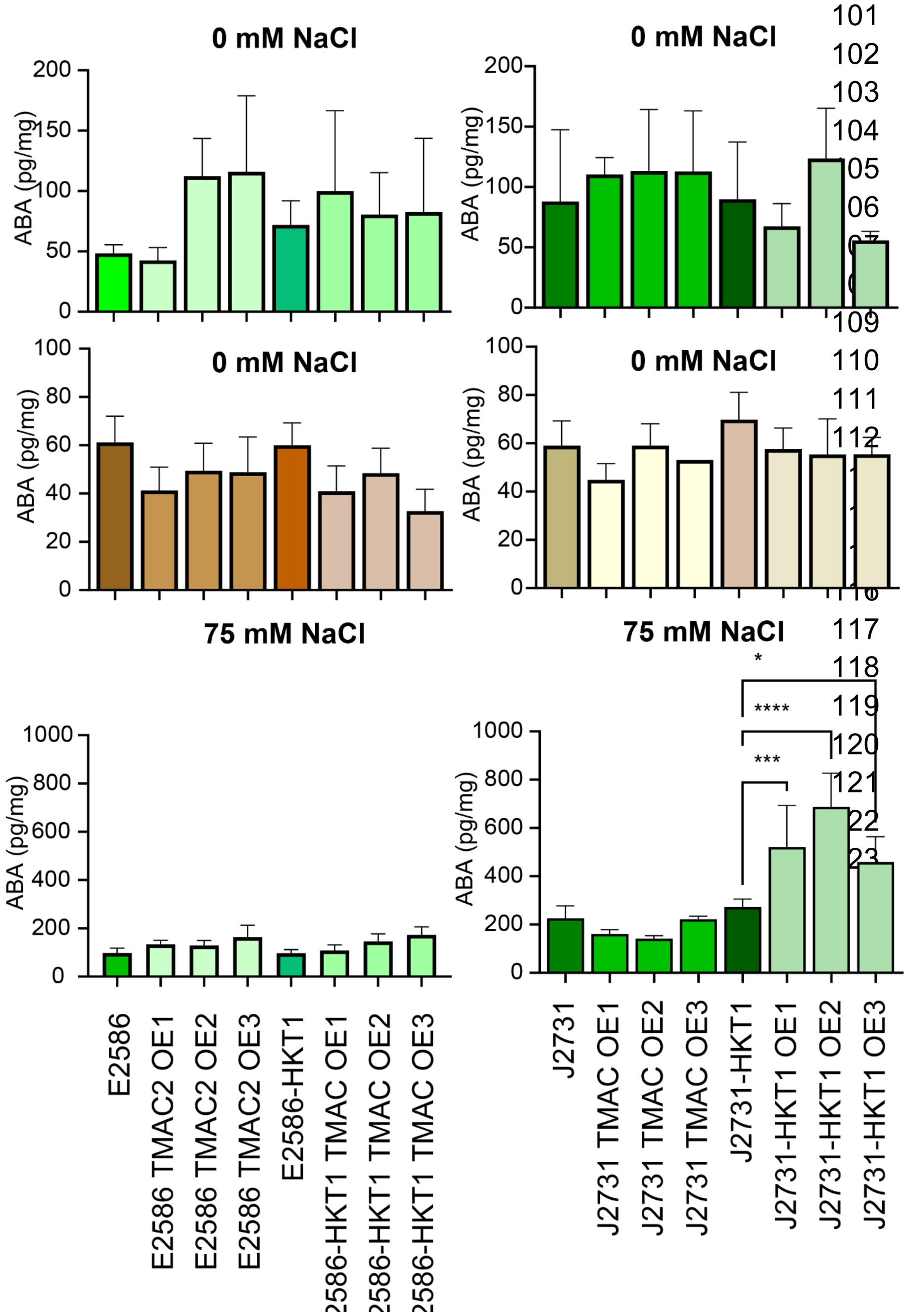


**Figure S5. Shoot tissue ABA accumulation in lines overexpressing TMAC2.** The ABA accumulation in Col-0 (E2586) and C24 (J2731) seedlings with and without tissue specific overexpression of HKT1 and additional over-expression of TMAC2 was measured in shoots (green graphs) and roots (brown graphs) Arabidopsis seedlings 21 days after transfer to 0 or 75 mM NaCl.

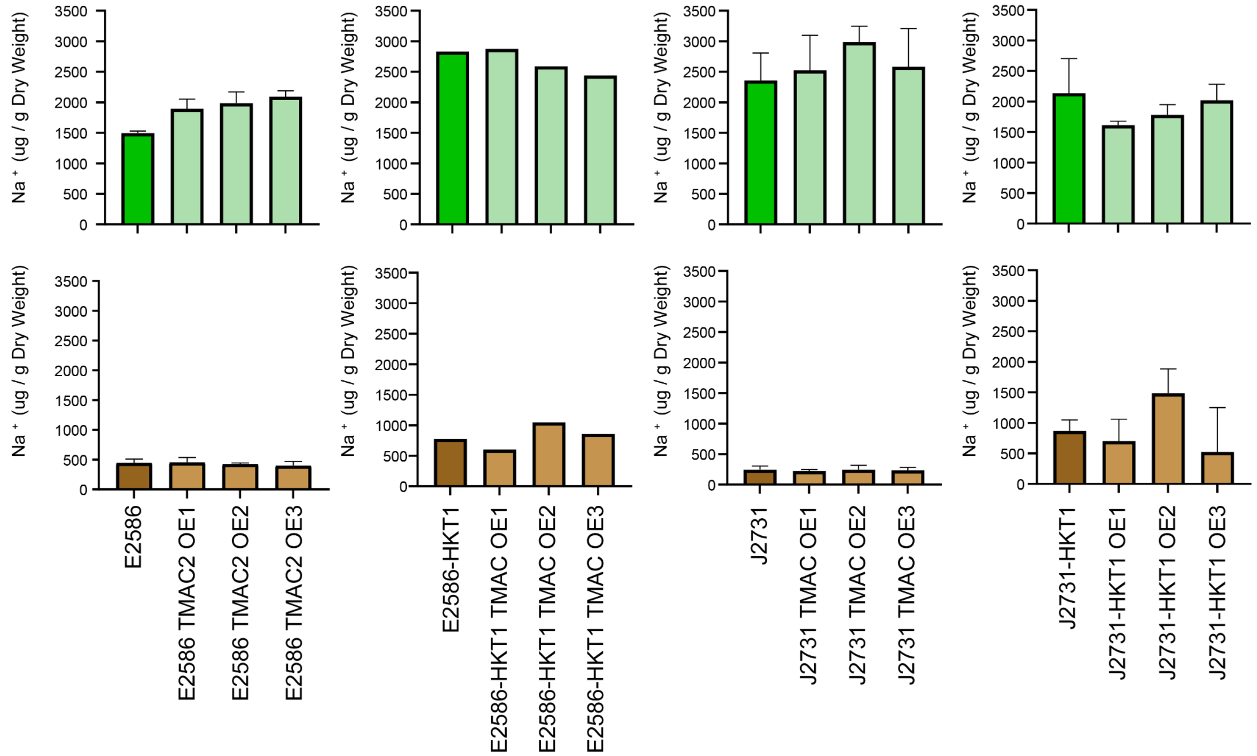


**Figure S6. Sodium ($Na^+$) accumulation in lines overexpressing TMAC2.** Four days old Arabidopsis seedlings from lines with tissue-specific HKT1 overexpression in Col-0 (E2586) and C24 (J2731) backgrounds with and without TMAC2 overexpression were exposed to salt stress (75 mM NaCl) for 21 days. The sodium ($Na^+$) accumulation was measured in seedling's shoot (green graphs) and root tissue (brown graphs). The bars represent the mean value calculated from at least 20 seedlings, and the error bars represent standard error. The significant differences between individual mutant lines and their respective background lines were determined using one-way ANOVA test, with *, **, *** and **** indicating p-values below 0.05, 0.01, 0.001 and 0.0001 respectively.

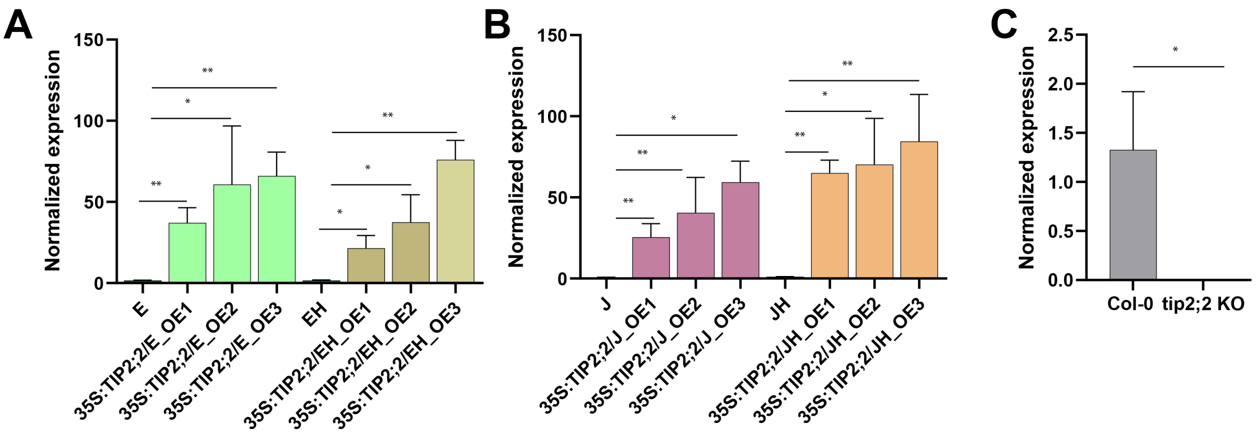


**Figure S7. Expression of TIP2;2 in generated overexpression lines.** Expression of TIP2;2 in Col-0 (E2586) and C24 (J2731) background with or without additional tissue-specific overexpression of HKT1. All expressions are based on three independent biological replicates collected from the leaves of soil grown plants. The significant differences between individual mutant lines and their respective background lines were determined using one-way ANOVA test, with *, **, *** and **** indicating p-values below 0.05, 0.01, 0.001 and 0.0001 respectively.

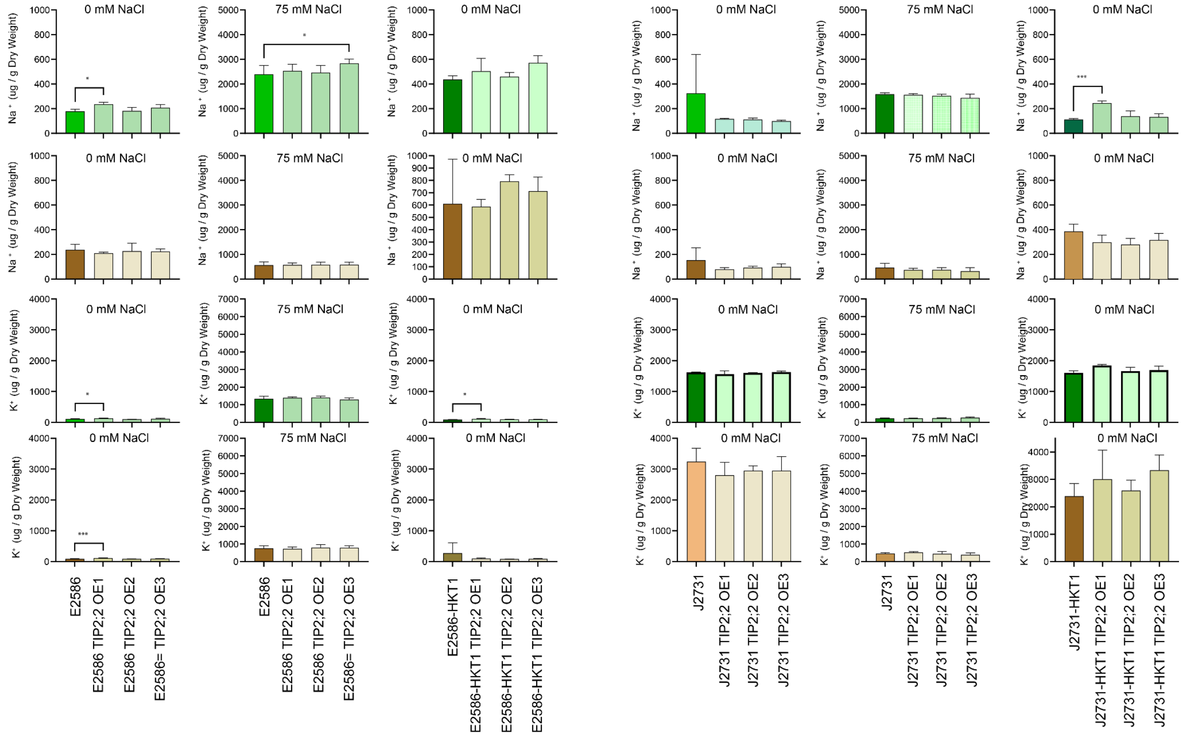


**Figure S8. Sodium (Na+) and potassium (K+) accumulation under control (0 mM NaCl) and salt (75 mM NaCl) conditions of TIP2;2 overexpressing lines.** Four days old Arabidopsis seedlings from lines with tissue-specific HKT1 overexpression in Col-0 (E2586) and C24 (J2731) backgrounds with and without TIP2;2 overexpression were exposed to salt stress (75 mM NaCl) for 21 days. The sodium ($Na^+$) and potassium ($K^+$) accumulation was measured in seedling's shoot (green graphs) and root tissue (brown graphs). The bars represent the mean value calculated from at least 20 seedlings, and the error bars represent standard error. The significant differences between individual mutant lines and their respective background lines were determined using one-way ANOVA test, with *, **, *** and **** indicating p-values below 0.05, 0.01, 0.001 and 0.0001 respectively.

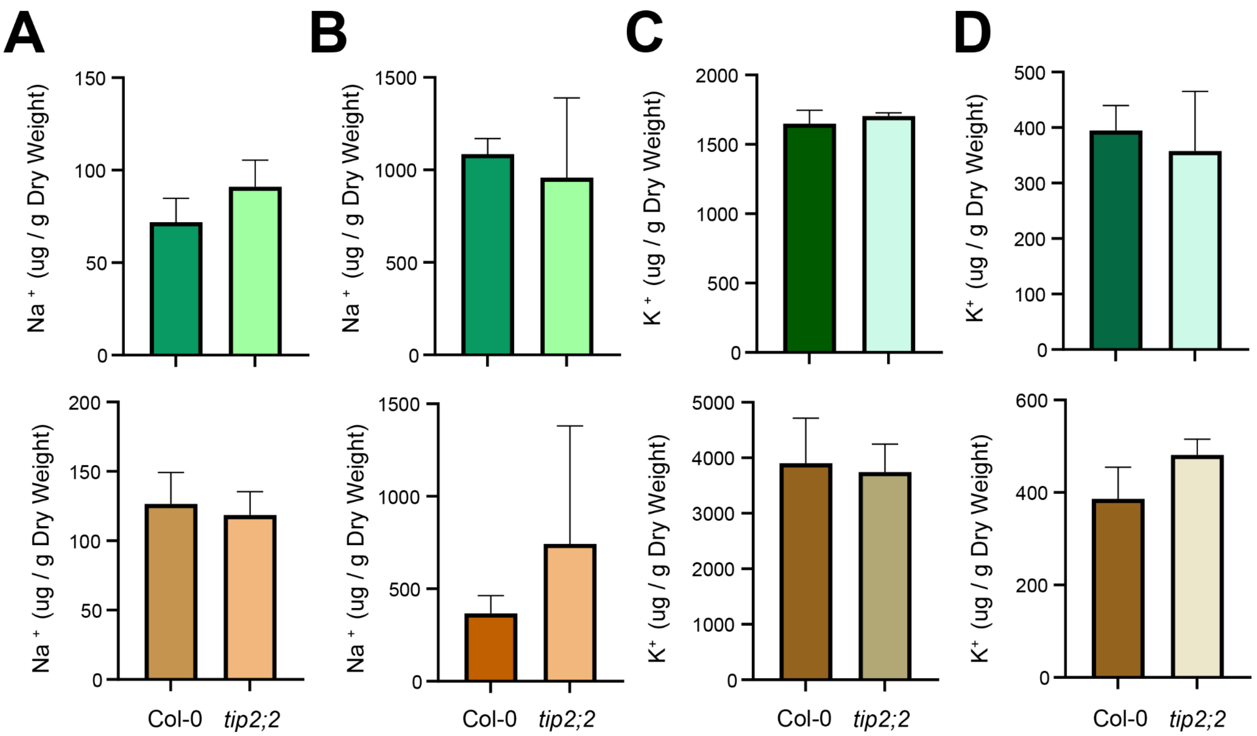


**Figure S9. Sodium ($Na^{+}$) and potassium ($K^{+}$) accumulation under control and salt stress conditions in tip2;2 mutant lines.** Four days old Arabidopsis seedlings from Col-0 and tip2;2 T-DNA insertion line (SALK_152463) were exposed to control (0 mM NaCl) or salt stress (75 mM NaCl) for 21 days. The sodium ($Na^{+}$) accumulation was measured in seedling's shoot (green graphs) and root tissue (brown graphs) under **(A)** control or **(B)** salt stress conditions. Additionally, potassium (K+) accumulation was also measured in seedling's shoot (green graphs) and root tissue (brown graphs) under **(C)** control or **(D)** salt stress conditions. The bars represent the mean value calculated from at least 20 seedlings, and the error bars represent standard error. The significant differences between individual mutant lines and their respective background lines were determined using one-way ANOVA test, with *, **, *** and **** indicating p-values below 0.05, 0.01, 0.001 and 0.0001 respectively.

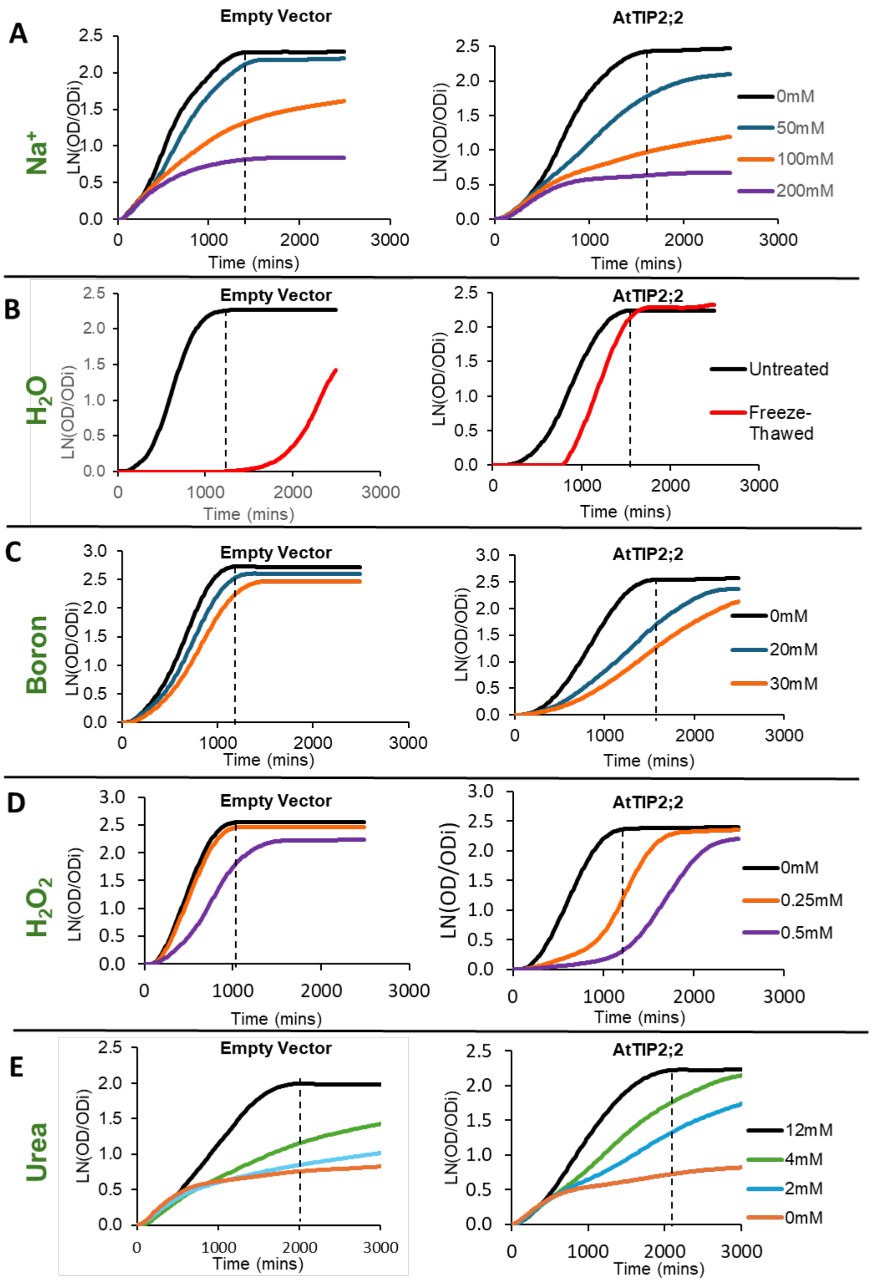


**Figure S10. Representative example yeast growth curves, Ln($OD/OD_i$) vs. time, of yeast expressing Empty vector Negative control and AtTIP2;2.** Respective yeast cultures were exposed to **(A)** 0mM, 50mM, 100MM and 200mM NaCl, for sodium toxicity assay; **(B)** Untreated and Freeze-thawed treated for Water transport assay; **(C)** 0mM, 20mM and 30mM Boric acid for Boron toxicity assay; **(D)** 0mM, 0.25mM and 0.5mM $H_2O2$, for $H_2O_2$ toxicity assay and **(E)** 12mM, 4mM, 2mM and 0mM Urea for urea growth-based screen. Growth comparisons shown in Figure 7 were captured from the area under the curves (AUC) until the vertical dashed lines (measuring time point) show on Ln($OD/OD_i$) vs. time graphs. AtTIP2;2- expressing yeast displays reduced growth over time at each of the sodium, boric acid, $H_2O_2$ concentrations, indicative of an increased sensitivity/toxicity response to these treatments compared to Empty vector control. AtTIP2;2-expressing yeast also shows increased survivorship/recovery post freeze-thaw treatment, compared to Empty vector control which shows minimal growth post treatment exposure. AtTIP2;2-expressing yeast showed enhanced yeast growth over time at low nitrogen media concentrations (2mM and 4mM urea) compared to Empty vector control

**Supplementary Table S1. List of mutants screened and primers used.**

**Supplementary Table S2. Primers used for cloning individual genes and links to their vector maps**

**Supplementary Table S3.List of primers used for expression**

**Supplementary Table S4. Relative expression of all mapped genes**

**Supplementary Table S5. Average and Standard Error of gene expression**
